# DNA methylation shows footprints of altitude selection in the clonal plant species *Fragaria vesca*

**DOI:** 10.1101/2024.03.19.585697

**Authors:** Audrey Le Veve, Iris Sammarco, Vít Latzel, Clément Lafon Placette

## Abstract

- Climate change threatens plant species, potentially pushing them beyond their adaptive capacities. DNA methylation and other epigenetic modifications may enable rapid adaptation to environmental changes by generating locally adapted phenotypes. These phenotypic changes can be inherited across generations and may become targets of natural selection. However, direct evidence for selection on epialleles remains scarce. Addressing this gap is crucial, as population survival may heavily rely on DNA methylation, especially in clonal plants with limited genetic diversity.
- We employed population genomics approaches to investigate altitude-driven selection on epigenetic sites in clonal offspring of natural woodland strawberry (*Fragaria vesca*) populations. These offspring were grown in a common garden and derived from seven populations spanning an altitudinal range.
- Our genomic, epigenomic, and transcriptomic analyses identified epialleles in clones exhibiting signs of selection related to altitude. These loci overlapped with genes involved in the O-methyltransferase activity, potentially aiding altitude adaptation through enhanced secondary metabolite production. Interestingly, these epialleles were mostly independent of genetic variation, suggesting they may have arisen stochastically or in response to environmental variation.
- These findings suggest that heritable epigenetic variation could help clonal species quickly adapt to environmental challenges as those related to varying altitudes and/or temperatures.

## Introduction

Given their sessile nature, plants need to constantly respond to changing environments, i.e. show phenotypic plasticity. Plants can adjust their phenotypes through epigenetic modifications, which involve mitotically (and partly meiotically) heritable changes in gene expression (Riggs and Porter, 1996; Lauria and Rossi, 2011 for a review in plants). Thus, some epigenetic marks can be stably inherited across generations (Miryeganeh and Saze, 2020; Verhoeven and Preite, 2014 for reviews), while others are reversible, changing dynamically in response to environmental stimuli (Feng, Jacobsen and Reik, 2010). In addition, some epigenetic marks are reset during sexual reproduction (Calarco et al., 2012; Ibarra et al., 2012; Wibowo et al., 2016), suggesting that bypassing this process through clonal propagation may allow for greater epigenetic inheritance. This enhanced epigenetic inheritance could be particularly important for the adaptation of clonal species, as it could equip them with increased phenotypic inheritance and compensate for their limited genetic polymorphism (Verhoeven and Preite, 2014 for review). If epigenetic marks are more stable in clonal species and yet more variable than genetic mutations, epigenetic variation might serve as a substrate for selection (van der Graaf et al., 2015), potentially compensating for their reduced reliance on genetic variation for long-term adaptation. This assumption remains, however, to be tested in natural populations.

Among all epigenetic marks, DNA methylation has received the most attention in evolutionary studies due to its high mitotic and meiotic stability, as it is a covalent modification of DNA (Riggs and Porter, 1996). In plants, DNA methylation occurs in three sequence contexts: CG, CHG and CHH (with H representing A, C or T; Finnegan et al., 1998), which play distinct roles (Niederhuth and Schmitz, 2017). In all sequence contexts, DNA methylation plays a pivotal role in repressing the mobilisation of transposable elements (TEs) (Zemach and Zilberman, 2010), while CG and CHH methylation alone appear to be connected to gene expression, however in a complex, non-linear and not fully understood fashion (Li et al,. 2012^a^; Gent et al,. 2013; Lang et al,. 2017; Rajkumar et al., 2020; Martin, Seymour and Gaut, 2021).

DNA methylation variation occurs at a considerably higher rate than genetic mutations (Becker et al,. 2011) and can arise in response to several factors, namely genetic, environmental and stochastic (Zhang et al., 2013; Dubin et al., 2015; Zhang et al., 2018; Johannes & Schmitz, 2019; Galanti et al., 2022; Rodríguez et al., 2022). Genetically determined DNA methylation variants stem from genetic modifications (Richards, 2006), while environmentally induced variants are solely influenced by environmental cues (Medrano et al., 2014; Kawakatsu et al., 2016). Finally, stochastic DNA methylation variants arise from errors in the maintenance of DNA methylation during DNA replication (Johannes and Schmitz, 2019 for review). Notably, both environmental and stochastic DNA methylation variants can potentially arise quickly in a population (Becker et al,. 2011; Zhang et al., 2013; Thiebaut et al., 2019). Recent studies have shed light on extensive natural epigenetic variation influenced by both genetic and environmental factors (Dubin et al., 2015; Kawakatsu et al., 2016; De Kort et al., 2020; Galanti et al., 2022; Sammarco et al., 2023) or exclusively by environmental conditions (Rodríguez et al., 2022). This environmental variation seemed to primarily affect CHH methylation, followed by CHG and CG (Rodríguez et al., 2022; Galanti et al., 2022; Sammarco et al., 2023). Some of these studies also revealed extensive inheritance of natural epigenetic variation across clonal generations, with the inheritance progressively decreasing from CG to CHG and CHH contexts (Wibowo et al., 2018; Rodríguez et al., 2022; Sammarco et al., 2024). While there is potential for these inherited variants to shape offspring phenotypes and facilitate rapid adaptation (Miryeganeh & Saze, 2020; Ashe et al., 2021), substantial empirical evidence remains elusive.

Thus, DNA methylation, as genetic mutations, has the potential to meet all the criteria required for natural selection to act: methylation profiles (or epi-alleles) can vary within and between natural populations (Schmitz et al., 2013^b^), be inherited across generations, and influence gene expression and, consequently, individual fitness (Miryeganeh and Saze, 2020). However, identifying selection on epigenomic regions in natural plant populations stays hard. A theoretical model demonstrated that selection on methylated sites is possible (Charlesworth and Jain, 2014). If this is true in natural conditions, this may provide novel insights on adaptive evolution (Charlesworth and Jain, 2014; Wang and Fan, 2015).

It is crucial to recognize that population epigenomics entails more than just screening for natural epigenetic diversity. According to the neutral theory of evolution, most observed (epi)genetic variation is not the result of selection. It is fundamental to distinguish the impact of selection from other factors affecting diversity in methylated sites, such as demographic history, mutation rates, genetic drift (Vitti, Grossman and Sabeti, 2013 for review), and environmental responses that are not necessarily adaptive. To achieve this, we can employ population genomics assumptions (Vitti, Grossman and Sabeti, 2013 for review of the assumptions; Wang and Fan, 2015 for a test on Arabidopsis and human genes), which predict that heritable epigenetic loci under selection should exhibit increased epi-differentiation between populations exposed to different selection pressures.

In this study, we investigate whether epigenetic variation is subject to natural selection in wild populations of the clonal plant *Fragaria vesca* (woodland strawberry). To address this, we employed a multi-step approach, using publicly available genomic, epigenomic, and transcriptomic data from clones originating from low- and high-altitude populations cultivated under common garden conditions (Sammarco et al. 2024). First, we employed classical population genomics metrics, specifically the fixation index (Fst), to identify heritable candidate regions under altitudinal selection. Next, we used an epigenome-wide association (EWA) approach to identify the epi-alleles associated with both altitude and phenotypic traits. Finally, we distinguished epi-alleles under selection that were regulated by genomic variation from those potentially arising through environmental or stochastic variation, revealing that most of the methylation profiles selected in the two altitude groups were non-genetically determined. By cultivating plants in a common garden, we minimised direct environmental effects, allowing us to focus on inherited epigenetic patterns and assess potential signatures of selection on epialleles in response to altitude.

## Materials and methods

### Study system

*Fragaria vesca* L. (*Rosaceae*) is a widespread herbaceous perennial species with a native range that includes Europe, northern Asia, North America, and northern Africa (Darrow, 1966). It reproduces both clonally through stolons and sexually through seeds, although sexual reproduction is very rare in natural conditions (Schulze et al., 2012). *F. vesca* is primarily a selfing species, but outcrossing is also possible (Li et al. 2012^b^; Hilmarsson et al., 2017).

### Plant collection and growth

The dataset studied was composed of seven populations of *F. vesca* from Italy from Sammarco et al. (2024), initially consisting of 21 natural populations from three European countries. The seven populations selected represented the largest altitudinal range (table S1). These populations were classified into two groups of altitudes, those situated at low altitudes (<1000m; three populations) and those situated at high altitudes (>1000m; four populations). With one exception, these populations also followed a temperature gradient, with low-altitude populations correlating with higher mean temperatures (>= 8°C), and vice versa (table S1). The exception was a high-altitude population (FV_IT_02) that was associated with lower temperatures.

Briefly, Sammarco et al. (2024) collected seven distinct individuals from field conditions from each of seven populations of *F. vesca* (n = 49) and planted them individually in the common garden of the Institute of Botany of the Czech Academy of Sciences in Průhonice, Czechia (49.994°N, 14.566°E; 323 m altitude). They let the plants propagate clonally for one year, and collected a fully developed leaf from offspring ramets of at least the third generation. They used these leaves for whole genome bisulfite sequencing (WGBS) analysis. From a subset of 21 plants in the common garden (3 plants per population), they also collected a fully developed leaf from each individual plant for RNA-sequencing.

### Methylation calling

We performed methylation calling separately for the three sequence contexts (CG, CHG and CHH), following the methods described in Sammarco et al. (2024). Specifically, the EpiDiverse WGBS pipeline (https://github.com/EpiDiverse/wgbs) was used for quality control, adaptor trimming, bisulfite reads mapping and methylation calling (Nunn et al., 2021). We used the most recent version of the *F. vesca* genome (v4.0.a2) in the mapping step (Edger et al., 2018; Li et al., 2019). All the cytosines having coverage ≥ 5 on the individual bedGraph files of methylated positions for each sample and sequence context were retained. For mapping statistics and bisulfite non-conversion rate, please refer to Sammarco et al. (2024).

### Conversion of methylation profiles into single nucleotide polymorphism(SNP)-like data

For each methylation context, the average methylation percentage by methylated position was calculated across all the samples to determine the methylation thresholds for categorising cytosines as methylated or unmethylated. The methylation averages were of 52.9, 23.2 and 4.1% for the CG, CHG and CHH contexts, respectively (table S2). The methylation thresholds were defined as the rounded values to the nearest whole number of these methylation averages, resulting in thresholds of 50%, 20%, and 4% for the CG, CHG, and CHH contexts, respectively. Based on these thresholds, the label “M” (indicating methylation) was assigned to sites exhibiting methylation percentages greater or equal to their corresponding threshold values. Conversely, the label “U” (indicating unmethylation) was assigned to cytosines exhibiting methylation percentages lower than their corresponding threshold value. Each individual was considered homozygous for either “U” or “M” alleles. Subsequently, the non-polymorphic methylation profiles were excluded. The summary of the number of cytosines considered is presented in table S2. The methylation profiles of each individual sample were converted in vcf format for polymorphism, Fst and linkage disequilibrium analyses (see below).

### Genomic data

Single nucleotide polymorphisms (SNPs) from WGBS data were inferred following the methods described in Sammarco et al. (2024). Specifically, the EpiDiverse SNP pipeline with default parameters (https://github.com/epidiverse/snp; Nunn et al., 2022) was used for each individual of all the populations grown under garden conditions (n=49). This pipeline was selected due to its high precision and sensitivity (Nunn et al., 2022). We then assessed the distribution of coverage for positions with a genotyping quality of 30 or higher to determine the mean coverage for 90% of the distribution (57 reads). Positions exceeding this mean coverage were excluded to minimise false SNP inference from repetitive elements. Finally, only sites covered by at least 3 reads in 80% of individuals from all populations were retained. The summary of the number of sites concerned is represented in table S2.

Then, using vcftools v0.1.15 (Danecek et al. 2011), we estimated the **π** on each site by population, the fixation index between populations of low and high altitude (Fst) and the linkage disequilibrium (R²) between all sites in overlapping windows of 30 kb in all the populations.

### Phenotypic measurements and analysis

For each plant, we recorded flower and fruit numbers as indicators of reproductive output. Additionally, aboveground biomass was estimated as a proxy for plant vigour, calculated by multiplying the total number of leaves by the length of the longest leaf, following Sammarco et al. (2022). We measured the length of the longest leaf (cm) on each plant as a key morphological trait. Stolon and ramet growth rates were also assessed to track vegetative propagation: their growth was measured weekly over a two-month period, with growth rates calculated by fitting an exponential growth model to the data using a log-linear regression approach. Stomatal density (number of stomata per mm²) and specific leaf area (SLA) were measured as indicators of physiological adaptation, following the methods described in Sammarco et al. (2023). The traits needed to follow a normal distribution, as this is a prerequisite for the linear mixed model (LMM) used in the association study (or GWA). After a Shapiro normality test, the distribution of continuous traits with a p-value lower than 0.05 was transformed into Log10. A principal component analysis (PCA) was then performed using the ggplot2 (version 2.2.1), factoextra (version 1.0.4) and FactoMiner (version 1.35) packages. A matrix representing the Spearman correlation coefficient (SC) between pairs of traits was constructed using the Hmisc (version 4.0.2) and corrplot (version 0.77) packages. After a hierarchical classification estimated by comparing the centred means of each trait in pairs, a correlation matrix of the traits between individuals was represented as a heatmap using the gplot package (version 3.0.1). Finally, the effect of the altitude of origin was evaluated using an ANOVA test and Tukey’s index (adjusted significance threshold padj=<0.05).

### Polymorphism of methylated sites between methylation context

Using vcftools v0.1.15 (Danecek et al. 2011), we calculated the polymorphism on methylated sites (**π***_met_*), separately for each sequence context and each population. Moreover, we distinguished between sites located in promoters, coding sequences and transposable elements (TEs). To test for significant differences in the **π***_met_* between methylation context and regions, we performed Kruskal– Wallis tests and Dunn’s test for multiple comparisons using rank sums, with two-sided P values adjusted using the Bonferroni method as implemented in the R package FSA (Ogle et al. 2020).

### Linkage disequilibrium of methylated sites

To assess the strength of the non-random association of alleles at different loci (linkage disequilibrium, LD), the coefficient of determination (R²) was calculated using vcftools v0.1.15 (Danecek et al. 2011). The R² values were calculated between variable methylated positions located in windows of 30kb and present in at least 80% of the individuals.

### Genetic and epigenetic structure and kinship

The Structure software 2.3.4 (Pritchard et al., 2000) was used to infer the number of ancestral populations based on 100,000 filtered SNPs or SMPs for Single Methylated Polymorphisms (randomly chosen between polymorphic sites in respective context) and thus to assign the 49 individuals to populations (Q matrix). The most likely number of clusters K in all simulations with admixture models was assumed to be in the range of K = 1 to K = 7. Ten replicates were conducted for each K with a burn-in period of 1000, followed by 500 MCMC steps. The ad hoc statistic ƊK was used to determine the most probable K (Evanno et al., 2005).

SPAGeDi software (Hardy and Vekemans, 2002) was used to estimate the Ritland (1995) matrix of pairwise kinship coefficients (K matrix) from the 100,000 filtered SNPs or SMPs using a 10,000 bootstrap resampling procedure.

### Detection of selection to altitude based on Fst of methylated sites

In population genomics, the fixation index (Fst) is typically used to detect selection and local adaptation. An increase in this index signifies a higher level of differentiation between populations, indicating directional selection. Using vcftools v0.1.15 (Danecek et al. 2011), we calculated the fixation index between methylated sites on populations of low and high altitude (Fst_met_), separately for each sequence context. Then, we defined the distribution of FST values for each context independently to determine the threshold of high FST as the FST value obtained at 99% of the total distribution. Finally, we defined the epigenomic regions candidate for selection in response to altitude as the regions of at least 140 bp (∼size of a nucleosome), including a minimum of 90% methylated sites separated by less than 1kb and with Fst met between populations from low and high altitude >= to the threshold.

To estimate the effects of the selection coefficient (S) and the forward and backward epimutation rates (***μ*** and ***ƞ***) on candidate regions for populations from low and high altitudes, we applied a method based on the analytical framework for hypermutable polymorphisms developed by Charlesworth and Jain (2014), implemented in the MCMCB model (http://rpubs.com/rossibarra/mcmcb; Xu et al,. 2020). First, we used mSFS from 1,000 randomly selected sites in the CG, CHG and CHH contexts from the populations at low and high altitude levels independently as control, alongside mSFS from candidate regions. Following description on Xu et al (2020), we estimated S, ***μ***, ***ƞ*** and the ratio ***μ***/***ƞ*** for both control and candidate mSFS using an effective population size (Ne) of 1,800,000 (Hilmarsson et al,. 2017). We ran the model using a chain length of N = 1,000,000 iterations, discarding the first 20% as burn-in.

### Genomic annotation of candidate regions for altitude-related selection

The overlaps of the candidate regions found by at least one of the method previously described with genes and TEs were assessed using bedtools v2.26.0 (Quinlan & Hall, 2010), with the v4.0.a2 gene annotations downloaded from the Genome Database for Rosaceae (GDR; https://www.rosaceae.org/species/fragaria_vesca/genome_v4.0.a2; Jung et al., 2019), and a TE annotation carried out using the EDTA annotation pipeline v1.9.6 (Ou et al., 2019) on the substituted genome using default parameters, kindly provided by López et al. (2022).

We conducted a Gene Ontology (GO) enrichment analysis on the candidate genes associated with altitude selection. This analysis was performed using the R package clusterProfiler (version 3.18.1; Yu et al., 2012). The results were considered statistically significant if they had a False Discovery Rate (FDR) adjusted P-value < 0.05.

### Detection of selection to altitude based on EWA analysis

We investigated methylation-phenotype relationships using an epigenome-wide association (EWA) analysis, employing a linear mixed model (MLM) as implemented in GEMMA (Xiang and Stephens, 2012), enabling the “-notsnp” option. The MLM is an extension of the classic linear model of the type *y=ax+b*, where *a* is a constant applied to a variable *x* and *b* is a starting constant. In this model, the constants *a* and *b* incorporate a set of fixed effects and random effects that account for the variability associated with each subject. In this study, the model explains the phenotypes by the sum of Q, K, the effect of each marker, and the residual, which accounts for all factors not captured by the model. We integrated the ten phenotypes previously mentioned and the altitude levels of origin of the populations as a binary phenotype (0 for low, 1 for high). The pvalues obtained were corrected using the FDR method with the qvalue package (version 2.30) to reduce the number of false positives linked to multiple tests. A statistical test of association was performed for each SNP/SMP and for each phenotype. To estimate the deviation of the pvalues from the MLM model, the distribution of the quantiles of the corrected pvalues as a function of the quantiles of the theoretical pvalues of each trait were represented in the form of qqplot, using the qqman package (versions 0.0.6 and 0.1.9 respectively). Finally, for the 11 phenotypes mentioned above, Manhattan plots were constructed using the same package, and a corresponding threshold Log10(FDR-corrected pvalues) = 1.3 was added to indicate significant associations.

Finally, to ensure the consistency of the analysis, we investigated genotype-phenotype relationships using a genome-wide association (GWA) analysis, employing the same linear mixed model implemented in GEMMA (Xiang and Stephens, 2012) after removing the “-notsnp” option.

### DMR calling and overlap with candidate regions

We identified differentially methylated regions (DMRs) using the EpiDiverse DMR pipeline (https://github.com/EpiDiverse/dmr) (Nunn et al., 2021) and the DMR caller *metilene* with default parameters (Jühling et al., 2016). For input, we used individual bedGraph files filtered to include only cytosines with a coverage of ≥ 5, and we called DMRs between low-altitude and high-altitude populations.

We examined the overlap between DMRs and methylation candidate regions, and tested for independence of the candidate regions from the DMRs with a X² test.

### Differential expression analysis and correlation analysis between methylation of candidate regions associated with altitude selection and DEGs

We analysed the RNA-sequencing data from Sammarco et al. (2024) to identify differentially expressed genes (DEGs) between populations from low and high altitudes. We performed the analysis using the DESeq2 package for R (v1.30.1; Love et al., 2014). Genes were considered differentially expressed if they had an adjusted P-value < 0.05 (Benjamini-Hochberg) and an absolute value of fold change (FC) ≥ 1.5.

To evaluate the correlation between the methylation of candidate regions linked to altitude selection and the expression of DEGs, we used the multi-block discriminant analysis DIABLO (Data Integration Analysis for Biomarker discovery using Latent cOmponents; Singh et al., 2019). DIABLO is a multivariate integrative classification method designed to identify correlated variables from heterogeneous datasets.

We ran the analysis separately for each sequence context, using the R package mixOmics (version 6.24.0; Rohart et al., 2017). We removed predictors with zero or near-zero variance, and we fine-tuned the number of components to retain in the model using the DIABLO’s perf function, with Mfold validation (n = 5) and cross-validation (nrepeat = 1000). The best performance was determined based on both the balanced error rate and centroid distances. We used methylation and gene expression of each individual sample (21 samples, 7 populations), and we used as predictive variables the altitude of the populations (high, low). We included all variables as these were part of a pre-selected group that showed differences in our predictive variables. This approach was feasible and efficient as the pre-selected group comprised a relatively low number of variables (30 DEGs, 72 candidate regions in CG, 58 in CHG and 2 in CHH). We built the final model using the block.splsda function designed to identify signatures of highly correlated variables across the multiple matrix sets (Singh et al., 2019). We visualised the correlations with the circosplot function.

### GWA analysis of genetic variants and methylation of candidate regions

To assess the putative genetic basis of altitude selection-associated candidate regions, we performed a genome-wide association (GWA) analysis. This approach originated from the assumption that if methylation patterns are genetically controlled, they should persist under garden conditions.

To increase the statistical robustness of the analysis, we included all the samples from Sammarco et al. (2024), encompassing populations from distinct climatic and geographic origins grown under garden conditions (21 populations, 7 plants per population; N = 147). Our rationale was that if a specific methylation pattern is under genetic control, this effect should be observable consistently across all populations, regardless of their climatic and geographic origins.

We conducted GWA analysis as outlined in Galanti et al. (2022), using the R package rrBLUP (4.6.1; Endelman, 2011). For genetic variants, we imputed the missing genotype calls using BEAGLE 5.2 (Browning et al., 2018) and filtered for minor allele frequency (MAF) > 0.04. After filtering, we retained 83,095 SNPs. To correct for population structure, we used an identity by state (IBS) matrix derived from variants filtered for MAF ≥ 0.01. These were also pruned for LD with an LD threshold (r²) of 0.8 for SNP pairs in a sliding window of 50 SNPs, sliding by 5.

For phenotypes, we employed individual average methylation across each candidate region. This was calculated using the regionCounts R function from the methylKit package (version 1.16.1; Akalin et al., 2012), with a minimum cytosine coverage of 5.

Finally, we selected the threshold P-value using Bonferroni correction to adjust for multiple comparisons.

## Results

### Phenotyping

We phenotyped plants in a common garden to assess whether those from different altitudes exhibited phenotypic variation. Only the longest leaf length differed significantly, with low-altitude plants having longer leaves than high-altitude plants (table S3; Fig. 1A). This trait was positively correlated with the biomass estimator and stomata number, and negatively correlated with the number of flowers, stolons and ramets (Fig 1B). The positive correlation between the biomass estimator and leaf length was expected, as the first one was calculated based on leaf dimensions.

**Fig 1:**
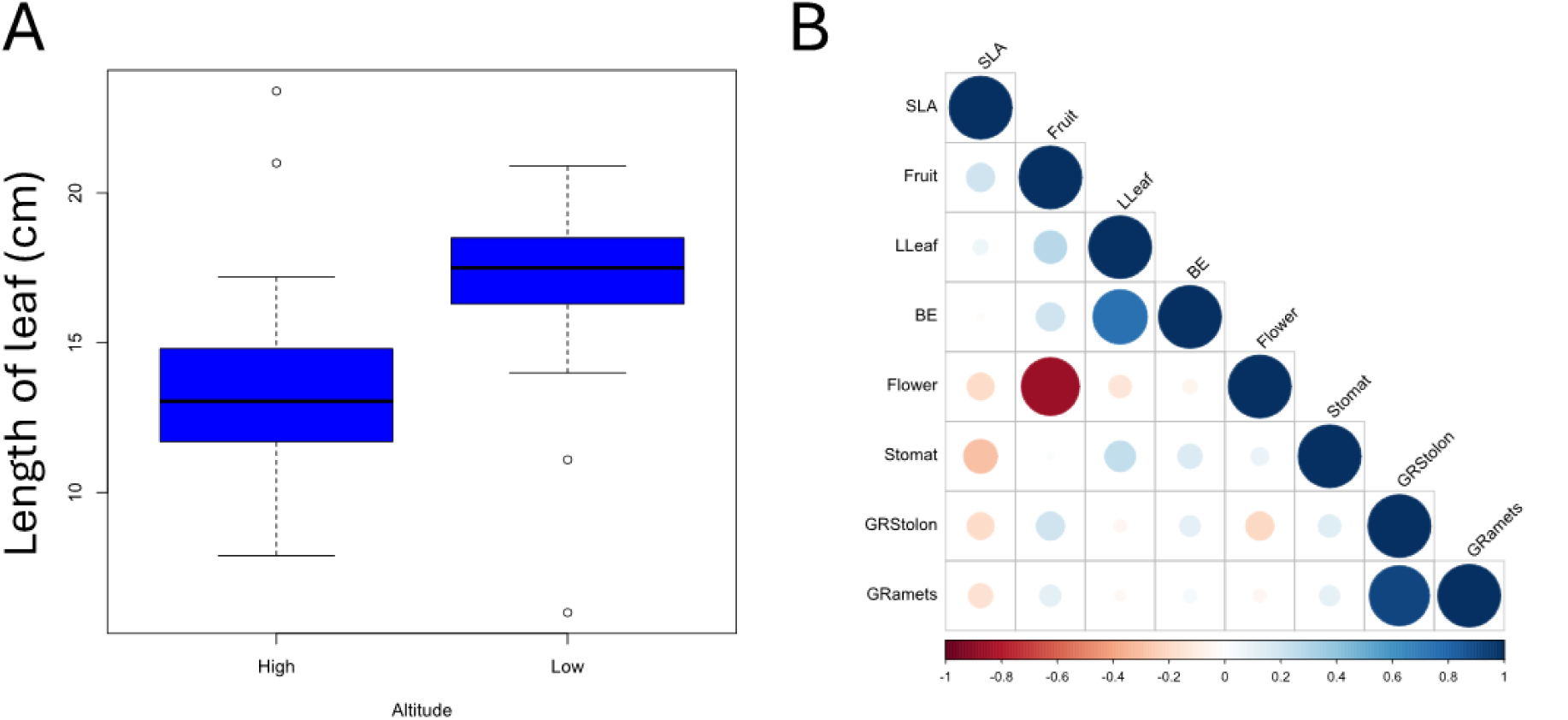
Variation of phenotypic traits across individuals from low- and high-altitude populations. A) Box-plot representations of the distribution for the length of the longest leaf (cm), which showed a significant association with altitude (Tukey’s test, P < 0.05). B) Lower matrix displaying the correlations between each analysed phenotype, with the Spearman correlation coefficient ranging from −1 (red) to +1 (blue). Stomat= mean number of stomata permm², Flower= number of flowers, SLA= Specific leaf area, Fruit= Number of fruits, GRStolon= stolons growth rate, GRramets= ramets growth rate, LLeaf= length of the longest leaf, BE= biomass estimate.

We then conducted a genome- and epigenome-wide association (G/EWA) analyses on this set of nine phenotypic traits but found no significant associations.

### Structuration of epigenome with altitude and demography

In order to characterize the effect of demography ald altitude levels on epigenome, we first compared their (epi)genetic polymorphism measured by **π** values using Kruskal-Wallis tests for sites overlapping transposable elements (TEs), the 5’ and 3’ untranslated regions (UTR), coding sequences, and promoters. Overall, we found no major differences in (epi)genetic polymorphism across altitude levels, regardless of the regions considered (table S4; Fig S1).

Population structure was apparent based on the ad hoc statistic ƊK, with the number of ancestral populations estimated to be one (K = 1) for the SNP and SMP datasets, except for the CHH sites, where K was estimated to be seven (K = 7). However, these seven groups did not exhibit clear trends explained by demographic factors (Fig S2).

To assess genetic and epigenetic relatedness among individuals and their association with altitude, we estimated pairwise kinship based on SNP and methylation data. SNP-based kinship estimates revealed a low degree of relatedness, with a mean of 0.0126, showing no correlation with altitude (Fig 2.A). In contrast, CG sites-relatedness was higher (mean = 0.0416) and correlated with altitude (Fig 2.B). CHG sites-relatedness also showed an altitude association but had a lower mean of 0.0072 (Fig 2.C). CHH sites-relatedness was minimal (mean = 0.0014) and uncorrelated with altitude (Fig 2.D).

**Fig 2:**
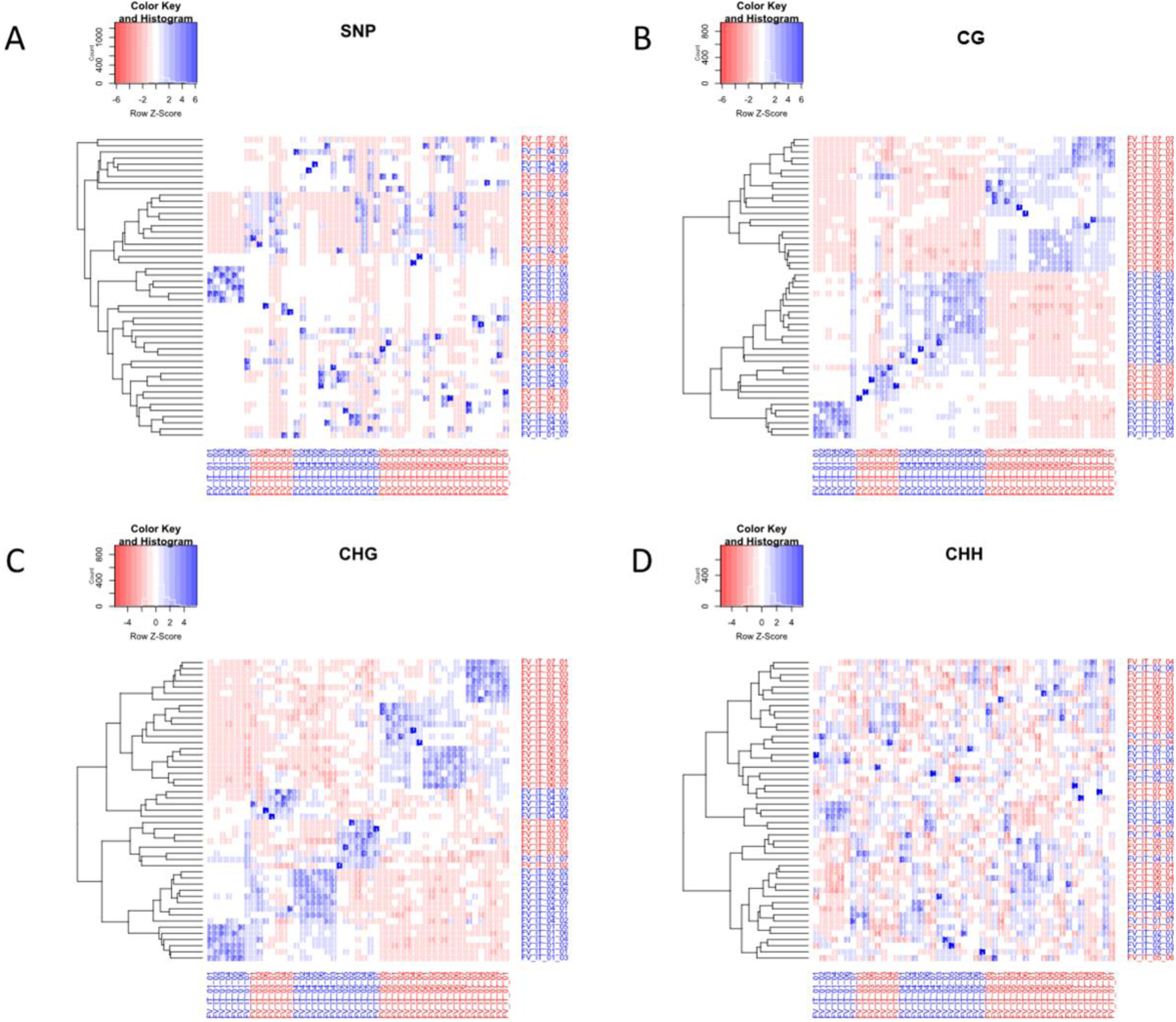
Kinship matrices among individuals based on genetic and methylation profiles. The matrices are derived from A) genotype (SNPs), B) CG, C) CHG, and D) CHH methylation profiles. The relatedness between individuals was estimated using SPAGEDI and is represented as a heatmap. Low normalised differentiations are represented in blue, while high normalised differentiations are shown in red. Hierarchical clustering is presented on the left. The colours assigned to the names indicate the different altitude clusters of origin, with red for > 1000m and blue for <1000m. N = 49 plants.

Kinship estimates from methylation data across all contexts were significantly and positively correlated with SNP-based estimates (Pearson’s p <2^e-16^ for all). However, the r² values were consistently below 0.21 (0.20, 0.07 and 0.02 for the CG, CHG and CHH context, respectively), suggesting a relatively low correlation between genetic and epigenetic relatedness.

Linkage disequilibrium (LD) can inform on the demographic history, diversity and effective recombination rates of populations. Pairwise LD estimates (R²) within genomic and methylation profiles revealed different levels of LD with distance (Fig 3). For genomic SNPs, the mean LD over distances of 30 kb ranged from 0.26 to 0.66. Specifically, the mean LD between two genomic SNPs remained relatively high, ranging from 0.27 to 0.37 for distances between 1 and 30,000 bp. In contrast, the mean LD between methylation sites was consistently low across all sequence contexts. For instance, the mean LD between two CG, CHG and CHH sites never exceeded 0.47, 0.37 and 0.02, respectively. The mean LD decreased substantially over the 30 kb range, by at least a factor of 10. The mean LD for variable sites separated by 1 and 30,000 bp was 0.46 and 0.03 for CG sites, 0.32 and 0.03 for CHG sites, and 0.10 and 0.03 for CHH sites. Notably, the mean LD between variable CG, CHG and CHH sites never exceeded 0.05 after distances of 2.5 kb, 13.5 kb and 9 bp, respectively.

**Fig 3:**
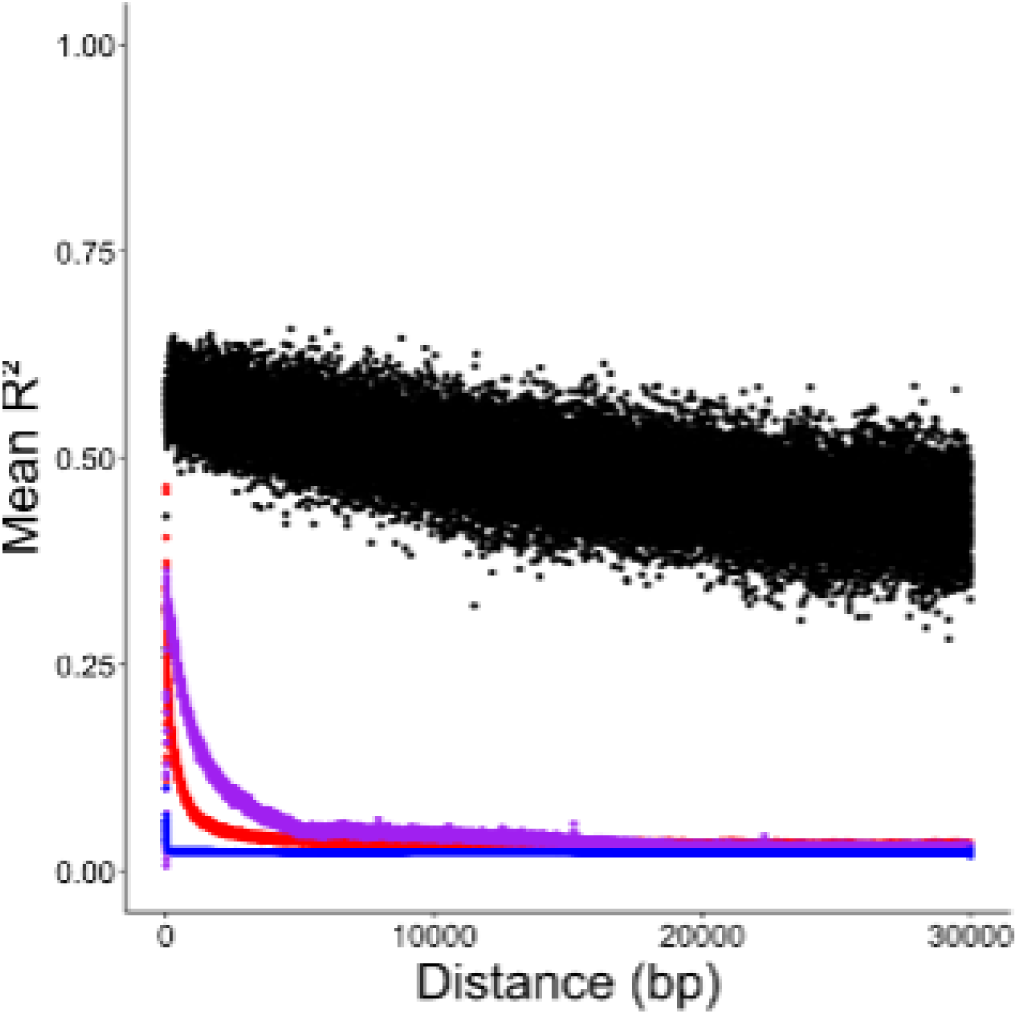
Decrease in linkage disequilibrium (LD) with distance between two variable sites across all methylation contexts. The mean R² values are presented based on the distance (in base pairs, bp) between two variable sites for CG (red), CHG (purple), CHH (blue) methylation contexts, and genetic SNPs (black). Only sites separated by less than 30 kb were considered.

### Candidate regions associated with selection by altitude

Classically, selection is detected by examining the decay of polymorphism against neutral expectations. However, the high epimutation rate modifies these assumptions, particularly when considering selection on epialleles (Charlesworth and Jain, 2014 and, Wang and Fan, 2015 for example on Tajima’s D). Therefore, we chose to detect selection driven by altitude by analysing polymorphism across populations with similar altitude ranges. Candidate regions associated with altitude were identified as those exceeding 140 bases (approximately the size of a nucleosome) and containing 90% or more sites with extreme FST values between populations from different altitudes, with sites spaced less than 1 kb apart. Extreme values of FST were defined as those falling within the top 1% of the distributions of all FST values for each genomic and epigenetic profile, assessed independently.

While no regions with significant genetic divergence were detected between low and high altitude populations using FST, we identified 72, 58, and 2 regions associated with altitude in the CG, CHG, and CHH methylation contexts, respectively (table S5). The candidate regions under selection in the CG and CHG context showed Gene Ontology (GO) enrichment related to O-methyltransferase activity (table S6). SMPs

Among the CG regions under selection for altitude, 13 (=18%) were not associated with any gene or TE, 13 (=18%) were located in the coding sequence of 14 genes, 50 (=69%) were located in the promoters of 58 genes and 8 (=12%) were associated with 9 TEs (table S5). For the CHG regions, we found 7 (=12%) not associated with any gene or TE, 14 (=24%) located in the coding sequence of 14 genes, 37 (=64%) were located in the promoters of 42 genes and 12 (=21%) associated with 14 TEs (table S5). In the CHH context, the 2 candidate regions under selection for altitude were located in the promoters of 3 genes (table S5).

We then assessed the effects of altitude on selection and epimutation rates in these candidate methylated profiles. we used the model MCMCBC based on mSFS (Charlesworth and Jain, 2014). First, we estimated the coefficient of selection S, the methylation epimutation rates (***μ***), the demethylation epimutation rates (***ƞ***) and the ratio ***μ***/***ƞ*** on mSFS based on 1,000 sites randomly chosen on epigenome as control regions in populations from low or high altitude levels. Regardless of the context, we found that the selection intensity (S) was multiplied by 7.34, 3.88 and 1.23 in CG, CHG and CHH context respectively in low-altitude compared to high-altitude populations (table S7), and we observed the same tendency for the ratio ***μ***/***ƞ*** (multiplied by 7.59, 17.6 and 1.66 in CG, CHG and CHH context respectively; table S7). Then, we used the median S and ***μ***/***ƞ*** obtained in the control positions in each population altitude to estimate the effect of these estimators on our candidate regions. First, as expected if altitude promotes selection on methylation profiles, all the candidate regions showed an S significantly different from their respective control regions (KW test, p < 0.05; table S8) and higher than in the control region (effect on S > 1; table S8) in one or both population altitudes, except for one candidate region in the CHG context on chromosome 2. Globally, these results confirm that the candidate regions found based on FST exhibited methylation profiles under selection.

To determine whether some SMPs were associated with altitude through a different approach, we conducted an epigenome-wide association (EWA) analysis using the percentage of methylation at each site (table S9). The altitude levels of origin were treated as a binary phenotype (0 = low altitude, 1 = high altitude). We retained the clustering of individuals identified with K=7 for the epigenome-wide association study (EWAS) on CHH profiles (Fig S2). For the other SMP datasets, we did not integrate population structure as a covariate.

For the CG sites, after false discovery rate (FDR) correction, 49 SMPs were found to be associated with altitude (Fig 4.A), but only one overlapped with a candidate region on chromosome 5 identified by FST analysis (Fig 5; table S9). Interestingly, this region also overlapped with one candidate region for the CHG context. For the CHG sites, 2,065 SMPs were associated with altitude after FDR correction (Fig 4.B), but none overlapped with the candidate regions identified by FST (table S9). We did not detect associations between methylation in the CG and CHG contexts and phenotypic traits. For the CHH sites, no associations were detected for any of the considered traits.

**Fig 4:**
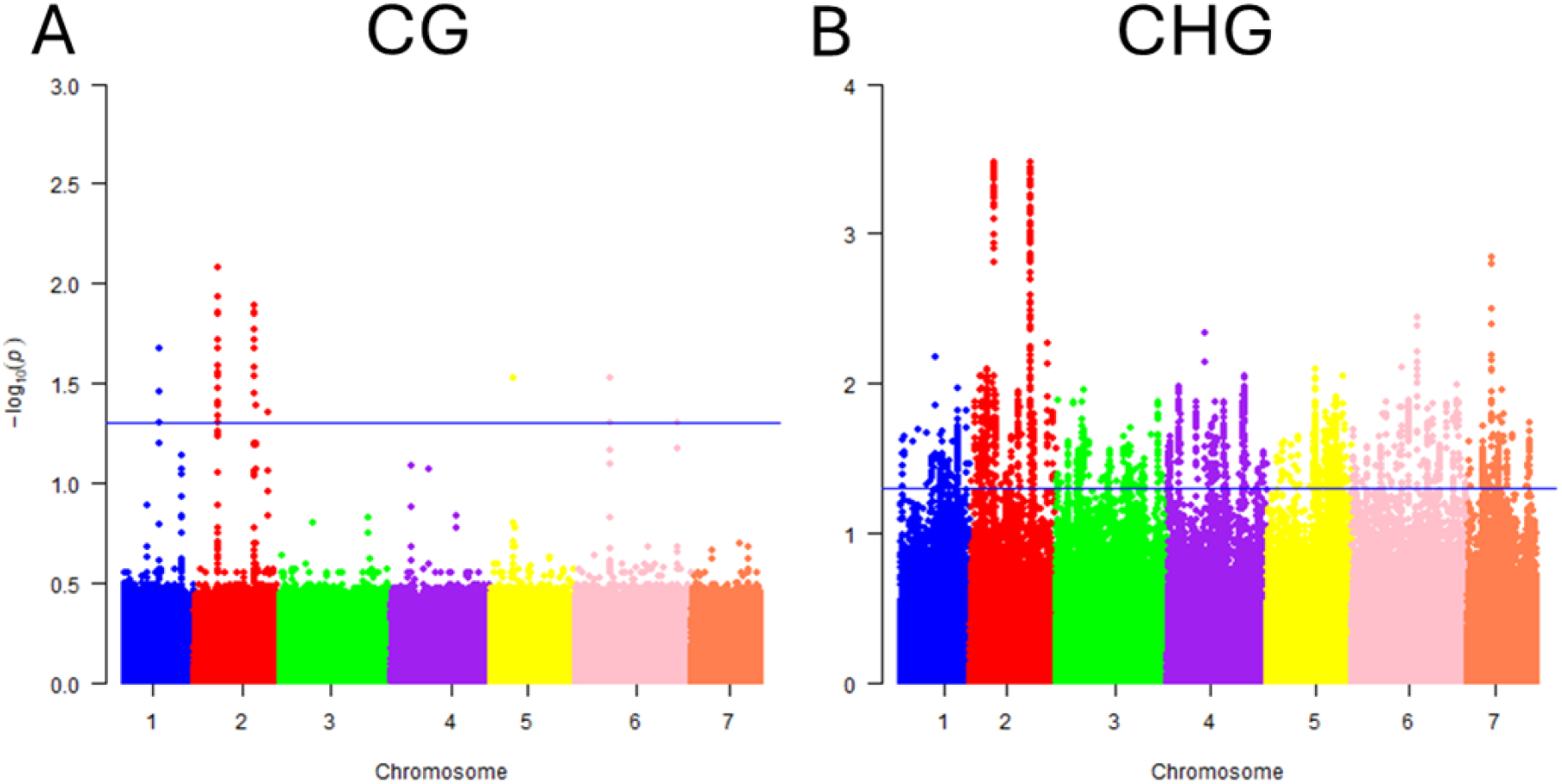
Epigenetic associations with altitude. Manhattan plots of epigenome-wide association (EWA) analyses for A) CG and B) CHG methylation contexts, displaying the 7 chromosomes (x-axis) and the – log10(P values) for each marker (y-axis). P values were obtained after false discovery rate (FDR) corrections. The blue solid line indicates the significance threshold (P value = 0.05).

**Fig 5:**
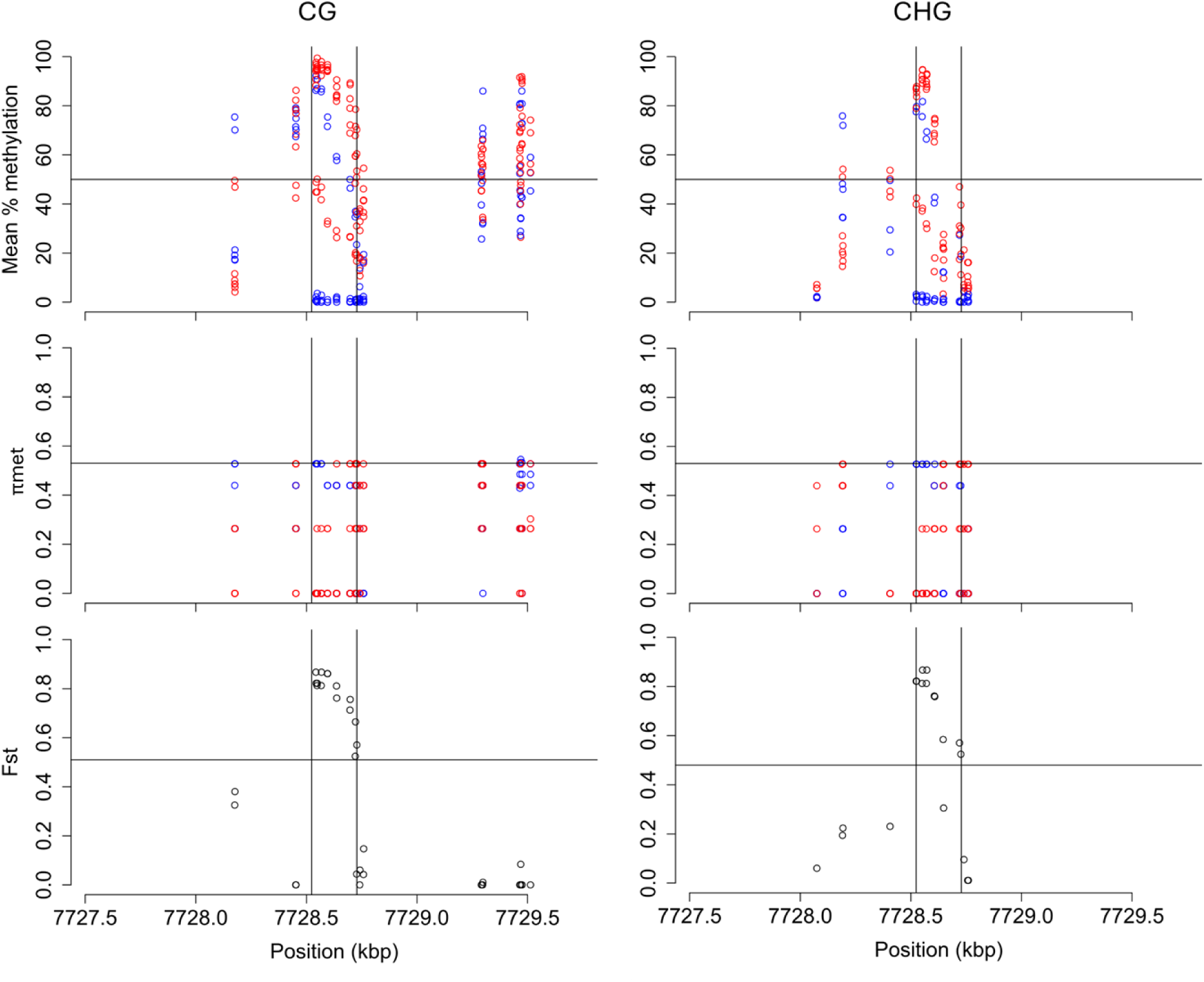
Variation in CG and CHG methylation profiles within a candidate region associated with altitude adaptation. The figure shows the variation in mean methylation levels (%), methylation diversity (**π**_met_), and Fst values between populations from different altitude levels for the CG (left) and CHG (right) contexts within a candidate region associated with altitude adaptation. Only variable sites were considered. The candidate region shown is located between positions 7,728,524 and 7,728,728 on chromosome 5. Horizontal black lines represent the respective thresholds for considering a position as methylated, based on the mean methylation levels, the **π**_met_ and the Fst values observed in 99% of the sites. The colours for the mean methylation levels and the **π**_met_ correspond to different altitude clusters, with red: > 1000m and blue: <1000m. N = 49 plants.

We compared our candidate regions with Differentially Methylated Regions (DMRs) to assess whether our candidate region-based approach yields results consistent with those obtained from DMRs, which are widely used in methylation studies. For the CG candidate regions, 32 overlapped with DMRs exhibiting hypermethylation in high-altitude populations, while an equal number overlapped with DMRs hypermethylated in low-altitude populations (43% overlap; table S5). For the CHG candidate regions, 37 overlapped with DMRs hypermethylated in high-altitude populations, compared to 19 hypermethylated in low-altitude populations (64 and 33% overlap, respectively; table S5). However, the CHH candidate regions did not overlap with any DMRs (table S5). The association between the candidate regions and the DMRs was statistically significant (X² test, p =1.11^e-12^).

We compared LD decay by distance (R²/bp) in the candidate regions overlapping with DMRs and 100 control regions of the same size as the median candidate region and DMR size, for each methylation context independently. For the CG context, the median size of the candidate regions and DMRs was 206 bp. The LD per base pair varied significantly across control regions, candidate regions, and DMRs (Kruskal-Wallis test, p= 2.10^e-14^), showing a significant decrease in DMRs compared to candidate and control regions (Dunn’s test, p= 9.57^e-09^ and 7.86^e-08^, respectively; Figure S3.A). In the CHG context, the median size of the candidate regions and DMRs was 259 bp. The LD per base pair also varied significantly across the three types of regions (Kruskal-Wallis test, p= 1.18^e-12^), with a significant increase in candidate regions and DMRs compared to controls (Dunn’s test, p= 1.12^e-08^ and 3.31^e-13^, respectively; Figure S3.B). For the CHH context, the median size of the candidate regions and DMRs was 58 bp. The LD per base pair varied significantly across the types of regions (Kruskal-Wallis test, p= 6.49^e-85^), with a significant increase in DMRs compared to controls (Dunn’s test, p= 4.55^e-86^; Figure S3.C).

### Genetic basis of candidate regions: Genome-Wide association (GWA) analysis

The methylation patterns of candidate regions under selection to altitude in the common garden may be influenced by underlying genetic factors. To investigate this, we conducted a GWA analysis using methylation levels of the candidate regions as phenotypes and SNPs as predictors (table S10).

For candidate regions in the CG, CHG, and CHH contexts, we detected 11, 8, and 0 significant associations, respectively, representing 15.3% of the candidate regions in the CG context, 13.8% in the CHG, and none in the CHH context. These significant associations involved a total of 57 SNPs, with 34 associated with candidate regions in the CG context and 23 in the CHG context. Notably, 16 SNPs (28.1%) were associated with at least two different candidate regions, regardless of the sequence context. Most SNPs overlapped with coding sequences (59% in the CG context and 75% in the CHG context), followed by gene promoters (18% in CG, 35% in CHG) and TEs (18% in CG, 17% in CHG).

Consequently, we found no significant associations for 61 regions in the CG context (85%), 50 in the CHG context (86%), and 2 in the CHH context (100%), suggesting that these regions are likely not under genetic control. The GO enrichment analysis for the CG context revealed enrichment of the term “O-methyltransferase activity,” which is crucial for lignin biosynthesis, stress tolerance, and disease resistance (Lam et al., 2007), while we found no significant GO enrichment for the CHG context.

### Correlation analysis between methylation of candidate regions associated with altitude selection and expression of DEGs

To assess the potential impact of methylation in candidate regions on gene expression in *cis* (i.e., where the candidate methylation regions overlap with the gene in which correlations are being tested), we overlapped the list of genes associated with these regions with those that were differentially expressed (DEGs) between low- and high-altitude populations. However, we found no overlap between the candidate regions and DEGs.

Next, we explored whether methylation of candidate regions might influence gene expression in *trans*, where the methylation region and the gene are not physically overlapping but still show correlated activity. Using the DIABLO method from mixOmics (Singh et al., 2019), we identified 54 total *trans* correlations in the CG, 40 in the CHG and 2 in the CHH context (Fig S5; table S11). These correlations were both positive (CG: 26, CHG: 31, CHH: 0) and negative (CG: 28, CHG: 9, CHH: 2), and implicated four genes related to chitin degradation, receptor-like kinase signalling, oxidation reduction activity and RNA-binding activity.

In the CG context, 32 candidate regions exhibited a significant *trans*-correlation with gene expression, representing 44% of all candidate regions. These genes were involved in phosphatidylinositol transfer, protein phosphatase activity, flavin-dependent monooxygenase activity, ribosomal protein synthesis, phosphate transport, and brassinosteroid response (table S11). In the CHG context, the 25 candidate regions (43%) correlated with gene expression included genes involved in histone methylation, protein folding, receptor-like kinase signalling, transcriptional regulation, and ribonuclease activity. In the CHH context,both candidate regions exhibited a significant *trans*-correlation with gene expression, with the associated genes involved in transcriptional regulation and lipid storage.

## Discussion

Just as changes in DNA sequence can create heritable genetic variation, epigenetic mechanisms, such as DNA methylation, can produce heritable epigenetic variants (epi-alleles) that influence gene expression and fitness (Miryeganeh et Saze, 2020). In clonal species, DNA methylation could be particularly essential for adaptation to environmental fluctuation (Verhoeven and Preite, 2014). However, although studies of whole epigenome diversity at single-nucleotide resolution exist, these types of analyses have mainly been applied to model organisms under controlled conditions (Cokus et al., 2008; He et al., 2010; Chodavarapu et al., 2012; Kawakatsu et al., 2016; Li, Varala, Moose, and Hudson, 2012; Schmitz et al., 2013^b^). Studies in natural populations are still mostly limited to low-resolution analysis and model species (Alonso et al., 2015; Schrey et al., 2013 for review on BS-AFLP). Here, we propose a comprehensive genomic, epigenomic and transcriptomic analysis at single-nucleotide resolution of clones of *F. vesca*, from natural populations sampled across an altitudinal gradient. Our objective was to identify epiloci under altitude-driven selection in a predominantly clonally reproducing plant species.

### The CG and the CHG contexts drive epigenetic differentiation in natural populations

In our study, we detected limited structure and variation in relatedness across individuals based on SMPs and SNPs. Consistent with these observations, we also found limited phenotypic variation among individuals from different altitude levels. Despite this weak (epi)genetic structure, individuals tended to cluster by population more distinctly based on the CG and CHG contexts than on genetic variation or CHH context. The epigenetic kinship showed only partial correlation with genetic kinship, suggesting that genomic factors alone did not fully account for the observed epigenetic variation. Thus, CG and CHG methylation patterns may reflect recent demographic history and could provide insights for reconstructing phyloepigenetic relationships (Sow et al., 2023). In contrast, in *A. thaliana*, only CG sites showed structure among wild populations, and this structure was largely explained by genomic factors (Schmitz et al., 2013^b^). Interestingly, the estimation of the coefficient of selection and epimutation rates on random sites suggests that, overall, the epigenome is more constrained for methylation in low-altitude populations than in high-altitude populations.

Although we studied DNA methylation in plants grown in a common garden, we also found that the epigenetic kinship in the CG and CHG contexts appeared to correspond with the altitude levels of origin. Altitude represents a broad environmental factor that influences more specific climatic variables, such as temperature. Our observations are consistent with multiple studies showing that DNA methylation patterns differ among populations in different natural environments (Lira-Medeiros et al., 2010; Richards et al., 2012; Medrano et al., 2014). Moreover, these variations can be linked to ecological variables and can occur independently of genetic differences. For example, Díez Rodríguez et al. (2022) found natural epigenetic variation in the clonal species *Populus nigra* linked to environmental variables and inherited across clonal generations. Therefore, our findings underscore the potential role of epigenetic variation in shaping adaptive responses in clonal species across diverse environmental gradients.

### The CHH context is the most dynamic methylation context

DNA methylation polymorphism (**π**_met_), which quantifies the epigenetic variation between individuals, showed an overall high level of polymorphism levels in SMPs compared to SNPs, specifically in the CHH context. This suggests a higher rate of epimutations in CHH compared to CG and CHG methylation in *F. vesca*. These results contrast with findings from studies in *A. thaliana*, which demonstrated that the CG context exhibits the most variability among individuals and generations grown under controlled laboratory conditions, while the CHG and CHH contexts display the lowest levels of spontaneous epimutation rates (Schmitz et al., 2011; Becker et al., 2011). This discrepancy may stem from the predominantly clonal reproduction of *F. vesca*. Vegetative propagation may lead to higher epimutation rates in CHH methylation compared to sexual reproduction, due to the absence of meiosis that resets many DNA methylation patterns, especially in this context (Feng et al., 2010; Calarco et al., 2012; Anastasiadi et al., 2021). This conclusion is supported by the extremely low linkage disequilibrium (LD) observed on CHH methylated sites, which is expected in a clonal species: in the absence of meiotic resetting, CHH methylation is more prone to gets easily (and randomly) lost through mitosis (Verhoeven and Preite, 2014 for review), reducing LD.

Furthermore, environmental variation is well recognized to directly impact DNA methylation patterns, especially in the CHH context. While the studied populations originated from low and high altitudes, the CHH kinship did not align with altitude, and very few CHH altitude candidate regions were identified compared to CG and CHG methylation. This discrepancy is likely due to the plants being grown in a common garden, where the new environment directly influenced CHH methylation patterns, exacerbated by the increased CHH epimutation rates associated with clonal propagation. Environmental effects may also explain why Schmitz et al. (2013b) found lower LD in the CHH context compared to CG and CHG, along with decreased conservation among natural populations of *A. thaliana* - findings that contrast with those of Schmitz et al. (2011) and Becker et al. (2011), who studied populations under highly controlled environments. Therefore, it is likely that environmental fluctuations play a major role in shaping DNA methylation diversity in nature, particularly in the CHH context, alongside reproductive strategies.

Modelling studies have predicted that an increase in mutation rates may provide short-term adaptive advantages to clonal organisms in changing environments (Sniegowski et al., 2000). In *A. thaliana*, Schmitz et al. (2011) estimated an epimutation rate of ∼4.46 x10^-4^ for CG sites per generation, while the genetic mutation rate is ∼7 ×10^−9^ per nucleotide per generation (Ossowski et al., 2010). This high frequency of epimutations suggests that epimutations can fastly uncouple from genetic mutations, especially in clonal species with reduced genetic diversity. Our LD estimates also support this, as LD remained high for genetic mutations but fastly decreased for epimutations (van der Graaf et al., 2015). While fast epimutation rates may offer short-term adaptive advantages, they could also limit long-term adaptability if the epimutations are too unstable or reversible (Kronholms & Collins, 2016). This may be the case for CHH epimutations, where we found fewer candidate regions with high FST values between low- and high-altitude populations. However, some CHH methylation variants persisted when the plants were transferred to the common garden, suggesting that these variants may be sufficiently stable to contribute to the adaptive potential of *F. vesca* populations.

### Selection of epigenomes and their interactions with genomes

For a trait to be subject to natural selection, it must affect an individual’s survival and/or reproductive success and be heritable (Darwin, 1859). In genetics, selection leads to frequency shifts favouring advantageous alleles, making these regions detectable (Pavlidis and Alachiotis, 2017 for review). This detectability is further enhanced as selection also affects the frequencies of neutral alleles linked to the selected ones (Smith and Haigh, 1974). Similar to genetic mutations, epimutations may significantly impact traits under selection, especially in clonal plants, which display a high degree of transgenerational conservation of methylation profiles (Miryeganeh and Saze, 2020; Verhoeven and Preite, 2014; for reviews on plant and clonal species, respectively). However, to our knowledge, no study has yet attempted to detect traces of selection on epigenomes. This is particularly noteworthy given that the survival of populations may be highly dependent on methylation, especially in the case of plants with limited genetic diversity, such as a clonal or highly inbred populations (Latzel et al., 2013; Latzel, Rendina González and Rosenthal, 2016).

In our study, we investigated DNA methylation Fst outliers among clones from populations across altitudinal gradients, grown in a common garden, to detect heritable regions under selection. High-altitude populations ranged from montane to subalpine zones (1,436-1,905 m), where environmental conditions, though not as extreme, represent mild selective pressures. The proportion of candidate regions conserved in the common garden, exhibiting significant Fst variation due to altitude of origin, varied among methylation contexts and often overlapped with differentially methylated regions (DMRs). However, overlap with SMPs significantly associated with altitude by EWAS was limited. The CG context presented the most regions with significant Fst variation, followed by CHG and CHH contexts, suggesting that CHH methylation patterns may be less relevant for natural selection. Consistent with this, we detected no association between SMPs in CHH and altitude of origin. Interestingly, among the 72 heritable CG candidate regions associated with altitude, 38% overlapped with candidate regions in the CHG context. This overlap suggests that adaptation through methylation likely involves coordinated CG and CHG methylation profiles. This coordination is supported by frequent associations of these candidate regions with genes sharing the same gene ontology term, specifically “O-methyltransferase activity”. O-methyltransferase activity is crucial for altitude adaptation, modulating secondary metabolites biosynthesis, including methylated flavonoids and phenols, which provide UV protection and antioxidant defence (Ali et al., 2024; Deng et al., 2024). Additionally, O-methyltransferases alter lignin composition, strengthening cell walls for mechanical resilience to high winds and reducing water loss through transpiration, critical under high-altitude conditions (e.g., extreme UV and limited water; Wang et al., 2018; Huang et al., 2024).

The genomic locations of these candidate methylation regions were primarily found in promoters, irrespective of the context considered. However, we observed no overlap between genes with differential expression associated with altitude and the candidate methylation regions, reflecting the complex and often context-dependent relationship between gene expressions and methylation (Dubin et al., 2015; Kawakatsu et al., 2016). Kawakatsu et al. (2016) proposed that the link between DNA methylation and gene expression might be overestimated and highly dependent on growth conditions. Moreover, methylation changes can also influence gene expression in *trans*, as we observed in our study, with both positive and negative correlations between methylation of candidate regions and gene expression, highlighting a complex interplay.

A key theoretical debate centres on whether selection can act on the epigenome independently of genomic variation. This possibility is particularly relevant for clonal species, which often have low genetic diversity (Verhoeven and Preite, 2014 for review). In our study, most of the 77 heritable candidate regions associated with altitude were not associated with genetic variants, suggesting that selection may act on methylation profiles without requiring genetic mutations. The limited number of associations observed between methylation of candidate regions and genetic variation may be due to the low genetic diversity within these populations, as evidenced by relatively few SNPs detected. Moreover, in this study we could not account for *de novo* TE insertions among the genetic variants. Given that these could be significant contributors to epigenetic variation, it is possible that we overlooked certain associations between genetic and DNA methylation variants. Methylation is particularly important for TE inactivation (Miryeganeh and Saze, 2020 for review), especially in clonally propagating populations. In these populations, TEs could quickly accumulate and lead to population extinction if not tightly repressed by DNA methylation (Charlesworth and Sniegowski, 1994; Dolgin and Charlesworth, 2006). Although a minority of genetic variants acting in *trans* on candidate methylated regions overlapped with TEs, TE polymorphisms, except in the CG context, were higher than in other genetic regions, suggesting relaxed selection.

## Conclusion

Our study on *Fragaria vesca* across an altitudinal gradient highlights the complex role of DNA methylation in natural populations, emphasising the differential impacts of CG, CHG and CHH contexts on epigenetic differentiation and adaptation potential. Our findings suggest that epigenetic variation can contribute to population differentiation and potentially aid adaptation in changing environments, even in the absence of genetic diversity. This underscores the importance of considering epigenetic mechanisms in ecological and evolutionary studies, especially for clonal species or those with limited genetic diversity. Furthermore, the intricate interplay between genetic and epigenetic factors in shaping phenotypic plasticity and adaptive potential reflects the complexity of natural selection in wild populations. Our study adds to a growing body of evidence suggesting that epigenetic mechanisms play a crucial role in plant adaptation, offering insights into the evolutionary significance of epigenetic variation in natural environments.

Speculatively, considering that many plant traits are controlled by multiple genes (polygenic control) and that epimutations are heritable, epigenetic variation could serve as a powerful means of generating phenotypic diversity without altering DNA sequence. This introduces the possibility of epigenetic mechanisms providing a dynamic and reversible layer of regulation that can rapidly respond to environmental changes, thus enhancing adaptation potential. In changing environments, such epigenetic flexibility, combined with polygenic control of traits, could increase a population’s evolutionary capacity, enabling the emergence of phenotypic diversity without relying solely on genetic mutations. For populations with low genetic variation, this adaptability could be crucial, offering an alternative and faster route to survival and reproductive success. Therefore, heritable epimutations may hold exciting potential for understanding the generation of biodiversity and the adaptability of organisms in response to environmental pressures.

## Acknowledgements

ALV and CLP have been supported by the Czech Science Foundation (GACR 22-20240S). IS and VL have been supported by the Czech Science Foundation (GACR 23-04749S) and by institutional research project RVO 67985939. IS has also been supported by a Postdoctoral Fellowship (PPLZ) awarded by the Czech Academy of Science.

Computational resources were supplied by the project ‘e-Infrastruktura CZ’ (e-INFRA CZ LM2018140) supported by the Ministry of Education, Youth and Sports of the Czech Republic.

## Author contributions

AL and IS developed and designed the experiments for the study. IS and VL performed the experiments. AL and IS analysed and interpreted the data. AL, IS and CL-P wrote the manuscript. All authors edited the manuscript.

## Supplementary tables list

Table S1: Environmental characteristics of the populations studied.

Table S2: Number of SNP and SMP analysed.

Table S3: Variation of phenotypic trait across populations of clones from different altitude levels

Table S4: Variation of mean **π** across populations from low and high altitude for the different genomic regions considered.

Table S5: Summary of the characteristics of each candidate region for selection by altitude.

Table S6: Gene Ontology (GO) enrichment analysis for candidate regions.

Table S7: SNPs significantly associated with candidate methylation regions and their genomic locations.

Table S8: Correlations between methylation of candidate regions and the expression of differentially expressed genes.

Table S9: SMPs significantly associated with phenotype or altitude level.

Table S10: SNPs significantly associated with candidate methylation regions and their genomic locations.

Table S11: Correlations between methylation of candidate regions and the expression of differentially expressed genes.

## Supplementaries figures list

**Fig S1:**
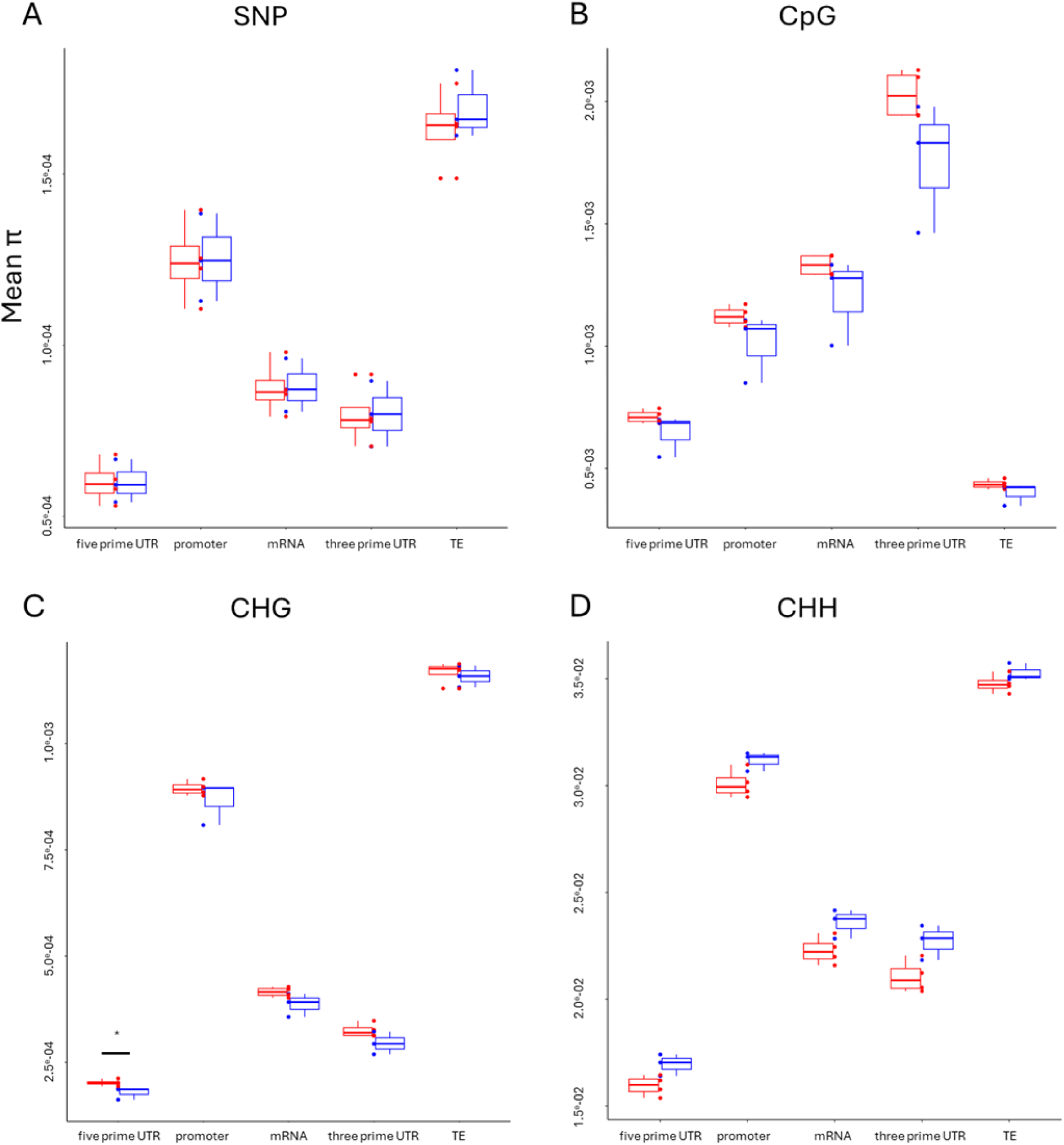
Mean **π** across populations from low and high altitude for the different genomic regions considered. Blue= populations from low altitude. red= populations from high altitude. *: p-values KW-test < 0.05. The non-significant p-values (>0.05) were not specified. N = 49 plants.

**Fig S2:**
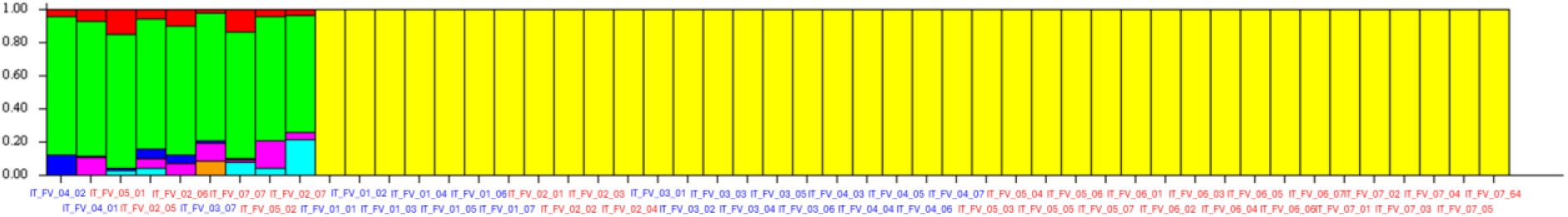
Structuration with k=7 for the CHH profiles. The names were sorted by Q and then by names. The colours assigned to the names represent the different altitude clusters in the field condition, with red: > 1000m and blue: <1000m. N = 49 plant

**Fig S3:**
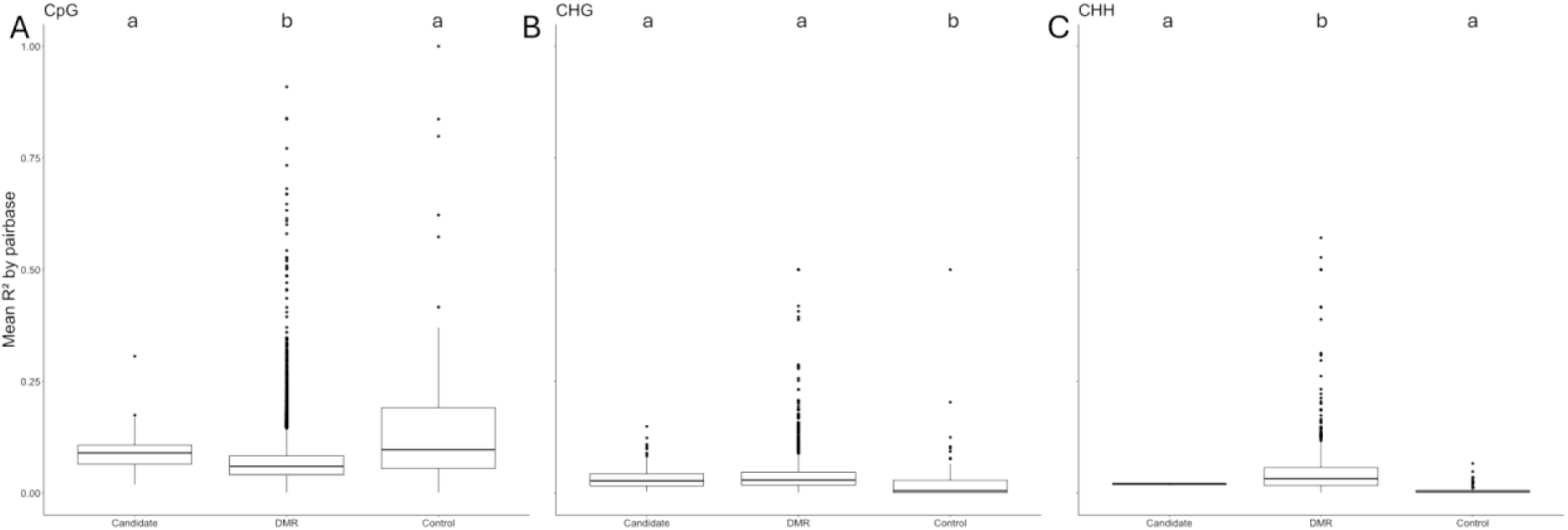
mean R² by base pairs for the candidate regions and the DMR compared to control regions for the three methylations profiles. The size of the control regions was defined as the median size found in candidate regions and DMR cumulated. The letters in lowercase signified the group significantly different according to Dunn’s test after Bonferroni’s correction (p-adj<0.05). N = 49 plants.

**Fig S4:**
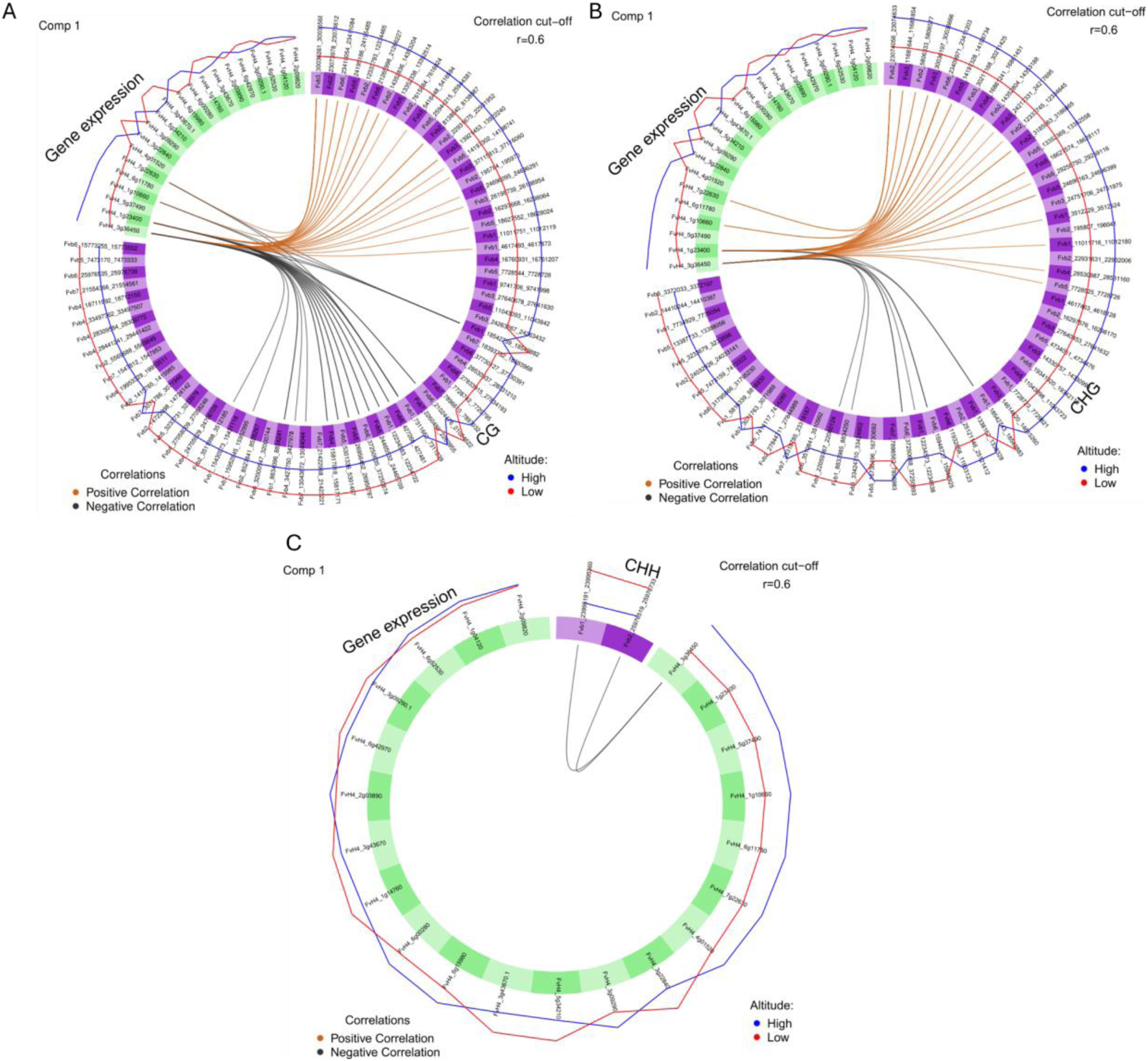
Circos plots showing correlations between methylation of candidate regions associated with altitude selection and the expression of differentially expressed genes. The different panels represent the methylation of candidate regions in the different sequence contexts: (a) CG, (b) CHG and (c) CHH. For each sequence context, we retained 1 component (Comp). The methylation dataset is depicted in purple, and the expression dataset in green. Within each Circos plot, the links represent the positive (orange) and negative (black) correlations, for which we used a correlation cutoff of r = 0.6. The blue and red lines outside the Circos plots correspond to the methylation and gene expression levels of the high and low altitude populations, respectively. These were utilised as predictive variables.

## References

Akalin, A., Kormaksson, M., Li, S., Garrett-Bakelman, F.E., Figueroa, M.E., Melnick, A., Mason, C.E., 2012. methylKit: a comprehensive R package for the analysis of genome-wide DNA methylation profiles. Genome Biology 13, R87. 10.1186/gb-2012-13-10-r87

Ali, A., Raza, A., Janiad, S., Rehman, S., Riaz, A., Khan, A., Deeba, F., I. Alalawy, A., Sakran, M., Alasmari, A., Moosa, A., Zulfiqar, F., 2024. Characterization of rice O-methyltransferase genes and their presumed homologs in Arabidopsis thaliana and Zea mays. Gene Reports 35, 101894. 10.1016/j.genrep.2024.101894

Alonso, C., Pérez, R., Bazaga, P., Herrera, C.M., 2015. Global DNA cytosine methylation as an evolving trait: phylogenetic signal and correlated evolution with genome size in angiosperms. Front Genet 6, 4. 10.3389/fgene.2015.00004

Anastasiadi, D., Venney, C.J., Bernatchez, L., Wellenreuther, M., 2021. Epigenetic inheritance and reproductive mode in plants and animals. Trends Ecol Evol 36, 1124–1140. 10.1016/j.tree.2021.08.006

Ashe, A., Colot, V., Oldroyd, B.P., 2021. How does epigenetics influence the course of evolution? Philosophical Transactions of the Royal Society B: Biological Sciences 376, 20200111. 10.1098/rstb.2020.0111

Atighi, M.R., Verstraeten, B., De Meyer, T., Kyndt, T., 2020. Genome-wide DNA hypomethylation shapes nematode pattern-triggered immunity in plants. New Phytologist 227, 545–558. 10.1111/nph.16532

Bates, D., Mächler, M., Bolker, B., Walker, S., 2015. Fitting Linear Mixed-Effects Models Using lme4. Journal of Statistical Software 67, 1–48. 10.18637/jss.v067.i01

Becker, C., Hagmann, J., Müller, J., Koenig, D., Stegle, O., Borgwardt, K., Weigel, D., 2011. Spontaneous epigenetic variation in the Arabidopsis thaliana methylome. Nature 480, 245–249. 10.1038/nature10555

Browning, B.L., Zhou, Y., Browning, S.R., 2018. A One-Penny Imputed Genome from Next-Generation Reference Panels. Am J Hum Genet 103, 338–348. 10.1016/j.ajhg.2018.07.015

Calarco, J.P., Borges, F., Donoghue, M.T.A., Van Ex, F., Jullien, P.E., Lopes, T., Gardner, R., Berger, F., Feijó, J.A., Becker, J.D., Martienssen, R.A., 2012. Reprogramming of DNA Methylation in Pollen Guides Epigenetic Inheritance via Small RNA. Cell 151, 194–205. 10.1016/j.cell.2012.09.001

Charlesworth, B., Jain, K., 2014. Purifying Selection, Drift, and Reversible Mutation with Arbitrarily High Mutation Rates. Genetics 198, 1587–1602. 10.1534/genetics.114.167973

Charlesworth, B., Sniegowski, P., Stephan, W., 1994. The evolutionary dynamics of repetitive DNA in eukaryotes. Nature 371, 215–220. 10.1038/371215a0

Chodavarapu, R.K., Feng, S., Ding, B., Simon, S.A., Lopez, D., Jia, Y., Wang, G.-L., Meyers, B.C., Jacobsen, S.E., Pellegrini, M., 2012. Transcriptome and methylome interactions in rice hybrids. Proceedings of the National Academy of Sciences 109, 12040–12045. 10.1073/pnas.1209297109

Cokus, S.J., Feng, S., Zhang, X., Chen, Z., Merriman, B., Haudenschild, C.D., Pradhan, S., Nelson, S.F., Pellegrini, M., Jacobsen, S.E., 2008. Shotgun bisulphite sequencing of the Arabidopsis genome reveals DNA methylation patterning. Nature 452, 215–219. 10.1038/nature06745

Danecek, P., Auton, A., Abecasis, G., Albers, C.A., Banks, E., DePristo, M.A., Handsaker, R.E., Lunter, G., Marth, G.T., Sherry, S.T., McVean, G., Durbin, R., 1000 Genomes Project Analysis Group, 2011. The variant call format and VCFtools. Bioinformatics 27, 2156–2158. 10.1093/bioinformatics/btr330

Darrow, G.M., 1966. The Strawberry. The Strawberry.

Darwin, C., 1859. On the Origin of Species. Harvard University Press.

De Kort, H., Panis, B., Deforce, D., Van Nieuwerburgh, F., Honnay, O., 2020. Ecological divergence of wild strawberry DNA methylation patterns at distinct spatial scales. Molecular Ecology 29, 4871– 4881. 10.1111/mec.15689

Deng, Y., Yang, P., Zhang, Q., Wu, Q., Feng, L., Shi, W., Peng, Q., Ding, L., Tan, X., Zhan, R., Ma, D., 2024. Genomic insights into the evolution of flavonoid biosynthesis and O-methyltransferase and glucosyltransferase in Chrysanthemum indicum. Cell Reports 43, 113725. 10.1016/j.celrep.2024.113725

Dolgin, E.S., Charlesworth, B., 2006. The Fate of Transposable Elements in Asexual Populations. Genetics 174, 817–827. 10.1534/genetics.106.060434

Dowen, R.H., Pelizzola, M., Schmitz, R.J., Lister, R., Dowen, J.M., Nery, J.R., Dixon, J.E., Ecker, J.R., 2012. Widespread dynamic DNA methylation in response to biotic stress. Proc Natl Acad Sci U S A 109, E2183–2191. 10.1073/pnas.1209329109

Dubin, M.J., Zhang, P., Meng, D., Remigereau, M.-S., Osborne, E.J., Paolo Casale, F., Drewe, P., Kahles, A., Jean, G., Vilhjálmsson, B., Jagoda, J., Irez, S., Voronin, V., Song, Q., Long, Q., Rätsch, G., Stegle, O., Clark, R.M., Nordborg, M., 2015. DNA methylation in Arabidopsis has a genetic basis and shows evidence of local adaptation. eLife 4, e05255. 10.7554/eLife.05255

Edger, P.P., VanBuren, R., Colle, M., Poorten, T.J., Wai, C.M., Niederhuth, C.E., Alger, E.I., Ou, S., Acharya, C.B., Wang, J., Callow, P., McKain, M.R., Shi, J., Collier, C., Xiong, Z., Mower, J.P., Slovin, J.P., Hytönen, T., Jiang, N., Childs, K.L., Knapp, S.J., 2018. Single-molecule sequencing and optical mapping yields an improved genome of woodland strawberry (Fragaria vesca) with chromosome-scale contiguity. GigaScience 7, gix124. 10.1093/gigascience/gix124

Endelman, JB. 2011. Ridge Regression and Other Kernels for Genomic Selection with R Package rrBLUP. The Plant Genome 4: 250–255.

Flanagan, B.A., Krueger-Hadfield, S.A., Murren, C.J., Nice, C.C., Strand, A.E., Sotka, E.E., 2021. Founder effects shape linkage disequilibrium and genomic diversity of a partially clonal invader. Molecular Ecology 30, 1962–1978. 10.1111/mec.15854

Feng, S., Jacobsen, S.E., Reik, W., 2010. Epigenetic reprogramming in plant and animal development. Science (New York, N.Y.) 330, 622. 10.1126/science.1190614

Finnegan, E.J., Genger, R.K., Peacock, W.J., Dennis, E.S., 1998. Dna Methylation in Plants. Annual Review of Plant Physiology and Plant Molecular Biology 49, 223–247. 10.1146/annurev.arplant.49.1.223

Galanti, D., Ramos-Cruz, D., Nunn, A., Rodríguez-Arévalo, I., Scheepens, J.F., Becker, C., Bossdorf, O., 2022. Genetic and environmental drivers of large-scale epigenetic variation in Thlaspi arvense. PLOS Genetics 18, e1010452. 10.1371/journal.pgen.1010452

Gent, J.I., Ellis, N.A., Guo, L., Harkess, A.E., Yao, Y., Zhang, X., Dawe, R.K., 2013. CHH islands: de novo DNA methylation in near-gene chromatin regulation in maize. Genome Res. 23, 628–637. 10.1101/gr.146985.112

He, G., Zhu, X., Elling, A.A., Chen, L., Wang, X., Guo, L., Liang, M., He, H., Zhang, H., Chen, F., Qi, Y., Chen, R., Deng, X.-W., 2010. Global Epigenetic and Transcriptional Trends among Two Rice Subspecies and Their Reciprocal Hybrids. The Plant Cell 22, 17–33. 10.1105/tpc.109.072041

Hilmarsson, H.S., Hytönen, T., Isobe, S., Göransson, M., Toivainen, T., Hallsson, J.H., 2017. Population genetic analysis of a global collection of Fragaria vesca using microsatellite markers. PLOS ONE 12, e0183384. 10.1371/journal.pone.0183384

Huang, E., Tang, J., Song, S., Yan, H., Yu, X., Luo, C., Chen, Y., Ji, H., Chen, A., Zhou, J., Liao, H., 2024. Caffeic acid O-methyltransferase from Ligusticum chuanxiong alleviates drought stress, and improves lignin and melatonin biosynthesis. Front. Plant Sci. 15. 10.3389/fpls.2024.1458296

Ibarra, C.A., Feng, X., Schoft, V.K., Hsieh, T.-F., Uzawa, R., Rodrigues, J.A., Zemach, A., Chumak, N., Machlicova, A., Nishimura, T., Rojas, D., Fischer, R.L., Tamaru, H., Zilberman, D., 2012. Active DNA Demethylation in Plant Companion Cells Reinforces Transposon Methylation in Gametes. Science 337, 1360–1364. 10.1126/science.1224839

Janko, K., Drozd, P., Eisner, J., 2011. Do clones degenerate over time? Explaining the genetic variability of asexuals through population genetic models. Biology Direct 6, 17. 10.1186/1745-6150-6-17

Johannes, F., Schmitz, R.J., 2019. Spontaneous epimutations in plants. New Phytologist 221, 1253– 1259. 10.1111/nph.15434

Jühling, F., Kretzmer, H., Bernhart, S.H., Otto, C., Stadler, P.F., Hoffmann, S., 2016. metilene: fast and sensitive calling of differentially methylated regions from bisulfite sequencing data. Genome Res 26, 256–262. 10.1101/gr.196394.115

Jung, S., Lee, T., Cheng, C.-H., Buble, K., Zheng, P., Yu, J., Humann, J., Ficklin, S.P., Gasic, K., Scott, K., Frank, M., Ru, S., Hough, H., Evans, K., Peace, C., Olmstead, M., DeVetter, L.W., McFerson, J., Coe, M., Wegrzyn, J.L., Staton, M.E., Abbott, A.G., Main, D., 2019. 15 years of GDR: New data and functionality in the Genome Database for Rosaceae. Nucleic Acids Research 47, D1137–D1145. 10.1093/nar/gky1000

Kawakatsu, T., Huang, S.C., Jupe, F., Sasaki, E., Schmitz, R.J., Urich, M.A. et al. 2016. Epigenomic Diversity in a Global Collection of Arabidopsis thaliana Accessions. Cell 166, 492–505. 10.1016/j.cell.2016.06.044

Kronholm, I., Collins, S., 2016. Epigenetic mutations can both help and hinder adaptive evolution. Molecular Ecology 25, 1856–1868. 10.1111/mec.13296

Lang, Z., Wang, Y., Tang, K., Tang, D., Datsenka, T., Cheng, J., Zhang, Y., Handa, A.K., Zhu, J.-K., 2017. Critical roles of DNA demethylation in the activation of ripening-induced genes and inhibition of ripening-repressed genes in tomato fruit. Proceedings of the National Academy of Sciences 114, E4511–E4519. 10.1073/pnas.1705233114

Latzel, V., Allan, E., Bortolini Silveira, A., Colot, V., Fischer, M., Bossdorf, O., 2013. Epigenetic diversity increases the productivity and stability of plant populations. Nat Commun 4, 2875. 10.1038/ncomms3875

Latzel, V., Rendina González, A.P., Rosenthal, J., 2016. Epigenetic Memory as a Basis for Intelligent Behavior in Clonal Plants. Frontiers in Plant Science 7.

Lauria, M., Rossi, V., 2011. Epigenetic control of gene regulation in plants. Biochimica et Biophysica Acta (BBA) - Gene Regulatory Mechanisms, Epigenetic control of cellular and developmental processes in plants 1809, 369–378. 10.1016/j.bbagrm.2011.03.002

Lam, K.C., Ibrahim, R.K., Behdad, B., Dayanandan, S., 2007. Structure, function, and evolution of plant O-methyltransferases. Genome 50, 1001–1013. 10.1139/G07-077

Li, X., Zhu, J., Hu, F., Ge, S., Ye, M., Xiang, H., Zhang, G., Zheng, X., Zhang, H., Zhang, S., Li, Q., Luo, R., Yu, C., Yu, J., Sun, J., Zou, X., Cao, X., Xie, X., Wang, J., Wang, W., 2012a. Single-base resolution maps of cultivated and wild rice methylomes and regulatory roles of DNA methylation in plant gene expression. BMC Genomics 13, 300. 10.1186/1471-2164-13-300

Li, J., Koski, M.H., Ashman, T.-L., 2012b. Functional characterization of gynodioecy in Fragaria vesca ssp. bracteata (Rosaceae). Annals of Botany 109, 545–552. 10.1093/aob/mcr279

Li, Y., Varala, K., Moose, S.P., Hudson, M.E., 2012. The Inheritance Pattern of 24 nt siRNA Clusters in Arabidopsis Hybrids Is Influenced by Proximity to Transposable Elements. PLOS ONE 7, e47043. 10.1371/journal.pone.0047043

Li, Y., Pi, M., Gao, Q., Liu, Z., Kang, C., 2019. Updated annotation of the wild strawberry Fragaria vesca V4 genome. Horticulture Research 6, 61. 10.1038/s41438-019-0142-6

Lira-Medeiros, C.F., Parisod, C., Fernandes, R.A., Mata, C.S., Cardoso, M.A., Ferreira, P.C.G., 2010. Epigenetic Variation in Mangrove Plants Occurring in Contrasting Natural Environment. PLOS ONE 5, e10326. 10.1371/journal.pone.0010326

López, M.-E., Roquis, D., Becker, C., Denoyes, B., Bucher, E., 2022. DNA methylation dynamics during stress response in woodland strawberry (Fragaria vesca). Horticulture Research 9, uhac174. 10.1093/hr/uhac174

Love, M.I., Huber, W., Anders, S., 2014. Moderated estimation of fold change and dispersion for RNA-seq data with DESeq2. Genome Biology 15, 550. 10.1186/s13059-014-0550-8

Martin, G.T., Seymour, D.K., Gaut, B.S., 2021. CHH Methylation Islands: A Nonconserved Feature of Grass Genomes That Is Positively Associated with Transposable Elements but Negatively Associated with Gene-Body Methylation. Genome Biology and Evolution 13, evab144. 10.1093/gbe/evab144

Medrano, M., Herrera, C.M., Bazaga, P., 2014. Epigenetic variation predicts regional and local intraspecific functional diversity in a perennial herb. Molecular Ecology 23, 4926–4938. 10.1111/mec.12911

Miryeganeh, M., Saze, H., 2020. Epigenetic inheritance and plant evolution. Population Ecology 62, 17–27. 10.1002/1438-390X.12018

Niederhuth, C.E., Schmitz, R.J., 2017. Putting DNA methylation in context: from genomes to gene expression in plants. Biochim Biophys Acta Gene Regul Mech 1860, 149–156. 10.1016/j.bbagrm.2016.08.009

Nunn, A., Can, S.N., Otto, C., Fasold, M., Díez Rodríguez, B., Fernández-Pozo, N., Rensing, S.A., Stadler, P.F., Langenberger, D., 2021. EpiDiverse Toolkit: a pipeline suite for the analysis of bisulfite sequencing data in ecological plant epigenetics. NAR Genomics and Bioinformatics 3, lqab106. 10.1093/nargab/lqab106

Nunn, A., Otto, C., Fasold, M., Stadler, P.F., Langenberger, D., 2022. Manipulating base quality scores enables variant calling from bisulfite sequencing alignments using conventional bayesian approaches. BMC Genomics 23, 477. 10.1186/s12864-022-08691-6

Ossowski, S., Schneeberger, K., Lucas-Lledó, J.I., Warthmann, N., Clark, R.M., Shaw, R.G., Weigel, D., Lynch, M., 2010. The Rate and Molecular Spectrum of Spontaneous Mutations in Arabidopsis thaliana. Science 327, 92–94. 10.1126/science.1180677

Ou, S., Su, W., Liao, Y., Chougule, K., Agda, J.R.A., Hellinga, A.J., Lugo, C.S.B., Elliott, T.A., Ware, D., Peterson, T., Jiang, N., Hirsch, C.N., Hufford, M.B., 2019. Benchmarking transposable element annotation methods for creation of a streamlined, comprehensive pipeline. Genome Biology 20, 275. 10.1186/s13059-019-1905-y

Pavlidis, P., Alachiotis, N., 2017. A survey of methods and tools to detect recent and strong positive selection. J of Biol Res-Thessaloniki 24, 7. 10.1186/s40709-017-0064-0

Qiao, Q., Edger, P.P., Xue, L., Qiong, L., Lu, J., Zhang, Y., Cao, Q. et al. 2021. Evolutionary history and pan-genome dynamics of strawberry (Fragaria spp.). Proc Natl Acad Sci U S A 118, e2105431118. 10.1073/pnas.2105431118

Quinlan, A.R., Hall, I.M., 2010. BEDTools: a flexible suite of utilities for comparing genomic features. Bioinformatics 26, 841–842. 10.1093/bioinformatics/btq033

Radersma, R., Noble, D.W.A., Uller, T., 2020. Plasticity leaves a phenotypic signature during local adaptation. Evolution Letters 4, 360–370. 10.1002/evl3.185

Rajkumar, M.S., Shankar, R., Garg, R., Jain, M., 2020. Bisulphite sequencing reveals dynamic DNA methylation under desiccation and salinity stresses in rice cultivars. Genomics 112, 3537–3548. 10.1016/j.ygeno.2020.04.005

Richards, E.J., 2006. Inherited epigenetic variation — revisiting soft inheritance. Nat Rev Genet 7, 395–401. 10.1038/nrg1834

Richards, C.L., Schrey, A.W., Pigliucci, M., 2012. Invasion of diverse habitats by few Japanese knotweed genotypes is correlated with epigenetic differentiation. Ecology Letters 15, 1016–1025. 10.1111/j.1461-0248.2012.01824.x

Riggs, A.D., Porter, T.N., 1996. Overview of Epigenetic Mechanisms. Cold Spring Harbor Monograph Archive 32, 29–45. 10.1101/0.29-45

Rodríguez, B.D., Galanti, D., Nunn, A., Peña-Ponton, C., Pérez-Bello, P., Sammarco, I., Jandrasits, K., Becker, C., Paoli, E.D., Verhoeven, K.J.F., Opgenoorth, L., Heer, K., 2022. Epigenetic variation in the Lombardy poplar along climatic gradients is independent of genetic structure and persists across clonal reproduction. 10.1101/2022.11.17.516862

Rohart, F., Gautier, B., Singh, A., Cao, K.-A.L., 2017. mixOmics: An R package for ‘omics feature selection and multiple data integration. PLOS Computational Biology 13, e1005752. 10.1371/journal.pcbi.1005752

Sammarco, I., Münzbergová, Z., Latzel, V., 2022. DNA Methylation Can Mediate Local Adaptation and Response to Climate Change in the Clonal Plant Fragaria vesca: Evidence From a European-Scale Reciprocal Transplant Experiment. Front. Plant Sci. 13. 10.3389/fpls.2022.827166

Sammarco, I., Münzbergová, Z., Latzel, V., 2023. Response of Fragaria vesca to projected change in temperature, water availability and concentration of CO2 in the atmosphere. Sci Rep 13, 10678. 10.1038/s41598-023-37901-8

Sammarco, I., Díez Rodríguez, B., Galanti, D., Nunn, A., Becker, C., Bossdorf, O., Münzbergová, Z., Latzel, V., 2024. DNA methylation in the wild: epigenetic transgenerational inheritance can mediate adaptation in clones of wild strawberry (Fragaria vesca). New Phytologist 241, 1621–1635. 10.1111/nph.19464

Schmitz, R.J., Schultz, M.D., Lewsey, M.G., O’Malley, R.C., Urich, M.A., Libiger, O., Schork, N.J., Ecker, J.R., 2011. Transgenerational Epigenetic Instability Is a Source of Novel Methylation Variants. Science 334, 369–373. 10.1126/science.1212959

Schmitz, R.J., He, Y., Valdés-López, O., Khan, S.M., Joshi, T., Urich, M.A., Nery, J.R., Diers, B., Xu, D., Stacey, G., Ecker, J.R., 2013a. Epigenome-wide inheritance of cytosine methylation variants in a recombinant inbred population. Genome Res. 23, 1663–1674. 10.1101/gr.152538.112

Schmitz, R.J., Schultz, M.D., Urich, M.A., Nery, J.R., Pelizzola, M., Libiger, O., Alix, A., McCosh, R.B., Chen, H., Schork, N.J., Ecker, J.R., 2013b. Patterns of population epigenomic diversity. Nature 495, 193–198. 10.1038/nature11968

Schrey, A.W., Alvarez, M., Foust, C.M., Kilvitis, H.J., Lee, J.D., Liebl, A.L., Martin, L.B., Richards, C.L., Robertson, M., 2013. Ecological Epigenetics: Beyond MS-AFLP. Integrative and Comparative Biology 53, 340–350. 10.1093/icb/ict012

Schulz, B., Eckstein, R.L., Durka, W., 2014. Epigenetic variation reflects dynamic habitat conditions in a rare floodplain herb. Molecular Ecology 23, 3523–3537. 10.1111/mec.12835

Schulze, J., Rufener, R., Erhardt, A., Stoll, P., 2012. The relative importance of sexual and clonal reproduction for population growth in the perennial herb Fragaria vesca. Popul Ecol 54, 369–380. 10.1007/s10144-012-0321-x

Singh, A., Shannon, C.P., Gautier, B., Rohart, F., Vacher, M., Tebbutt, S.J., Lê Cao, K.-A., 2019. DIABLO: an integrative approach for identifying key molecular drivers from multi-omics assays. Bioinformatics 35, 3055–3062. 10.1093/bioinformatics/bty1054

Smith, J.M., Haigh, J., 1974. The hitch-hiking effect of a favourable gene. Genetics Research 23, 23–35. 10.1017/S0016672300014634

Sniegowski, P.D., Gerrish, P.J., Johnson, T., Shaver, A., 2000. The evolution of mutation rates: separating causes from consequences. BioEssays 22, 1057–1066. 10.1002/1521-1878(200012)22:12

Sow, M.D., Rogier, O., Lesur, I., Daviaud, C., Mardoc, E., Sanou, E., Duvaux, L., Civan, P., Delaunay, A., Descauses, M.-C.L.-, Benoit, V., Le-Jan, I., Buret, C., Besse, C., Durufle, H., Fichot, R., Le-Provost, G., Guichoux, E., Boury, C., Garnier, A., Senhaji-Rachik, A., Jorge, V., Ambroise, C., Tost, J., Plomion, C., Segura, V., Maury, S., Salse, J., 2023. Epigenetic Variation in Tree Evolution: a case study in black poplar (Populus nigra). 10.1101/2023.07.16.549253

Thiebaut, F., Hemerly, A.S., Ferreira, P.C.G., 2019. A Role for Epigenetic Regulation in the Adaptation and Stress Responses of Non-model Plants. Frontiers in Plant Science 10. 10.3389/fpls.2019.00246

van der Graaf, A., Wardenaar, R., Neumann, D.A., Taudt, A., Shaw, R.G., Jansen, R.C., Schmitz, R.J., Colomé-Tatché, M., Johannes, F., 2015. Rate, spectrum, and evolutionary dynamics of spontaneous epimutations. Proceedings of the National Academy of Sciences 112, 6676–6681. 10.1073/pnas.1424254112

Verhoeven, K.J.F., Preite, V., 2014. Epigenetic variation in asexually reproducing organisms. Evolution 68, 644–655. 10.1111/evo.12320

Vitti, J.J., Grossman, S.R., Sabeti, P.C., 2013. Detecting Natural Selection in Genomic Data. Annual Review of Genetics 47, 97–120. 10.1146/annurev-genet-111212-133526

Wang, J., Fan, C., 2015. A Neutrality Test for Detecting Selection on DNA Methylation Using Single Methylation Polymorphism Frequency Spectrum. Genome Biology and Evolution 7, 154–171. 10.1093/gbe/

Wang, M., Zhu, X., Wang, K., Lu, C., Luo, M., Shan, T., Zhang, Z., 2018. A wheat caffeic acid 3-O-methyltransferase TaCOMT-3D positively contributes to both resistance to sharp eyespot disease and stem mechanical strength. Sci Rep 8, 6543. 10.1038/s41598-018-24884-0

Whitlock, M.C., 2008. Evolutionary inference from QST. Molecular Ecology 17, 1885–1896. 10.1111/j.1365-294X.2008.03712.x

Wibowo, A., Becker, C., Marconi, G., Durr, J., Price, J., Hagmann, J., Papareddy, R., Putra, H., SMPKageyama, J., Becker, J., Weigel, D., Gutierrez-Marcos, J., 2016. Hyperosmotic stress memory in Arabidopsis is mediated by distinct epigenetically labile sites in the genome and is restricted in the male germline by DNA glycosylase activity. eLife 5, e13546. 10.7554/eLife.13546

Wibowo, A., Becker, C., Durr, J., Price, J., Spaepen, S., Hilton, S., Putra, H., Papareddy, R., Saintain, Q., Harvey, S., Bending, G.D., Schulze-Lefert, P., Weigel, D., Gutierrez-Marcos, J., 2018. Partial maintenance of organ-specific epigenetic marks during plant asexual reproduction leads to heritable phenotypic variation. Proceedings of the National Academy of Sciences 115, E9145–E9152. 10.1073/pnas.1805371115

Wood, D.P., Holmberg, J.A., Osborne, O.G., Helmstetter, A.J., Dunning, L.T., Ellison, A.R., Smith, R.J., Lighten, J., Papadopulos, A.S.T., 2023. Genetic assimilation of ancestral plasticity during parallel adaptation to zinc contamination in Silene uniflora. Nat Ecol Evol 7, 414–423. 10.1038/s41559-022-01975-w

Xu, G., Lyu, J., Li, Q., Liu, H., Wang, D., Zhang, M., Springer, N.M., Ross-Ibarra, J., Yang, J., 2020. Evolutionary and functional genomics of DNA methylation in maize domestication and improvement. Nat Commun 11, 5539. 10.1038/s41467-020-19333-4

Yu, G., Wang, L.-G., Han, Y., He, Q.-Y., 2012. clusterProfiler: an R Package for Comparing Biological Themes Among Gene Clusters. OMICS: A Journal of Integrative Biology 16, 284–287. 10.1089/omi.2011.0118

Zhang, Y.-Y., Fischer, M., Colot, V., Bossdorf, O., 2013. Epigenetic variation creates potential for evolution of plant phenotypic plasticity. New Phytologist 197, 314–322. 10.1111/nph.12010

Zhang, Q., Liang, Z., Cui, X., Ji, C., Li, Y., Zhang, P., Liu, J., Riaz, A., Yao, P., Liu, M., Wang, Y., Lu, T., Yu, H., Yang, D., Zheng, H., Gu, X., 2018. N6-Methyladenine DNA Methylation in Japonica and Indica Rice Genomes and Its Association with Gene Expression, Plant Development, and Stress Responses. Molecular Plant 11, 1492–1508. 10.1016/j.molp.2018.11.005

Zemach, A., Zilberman, D., 2010. Evolution of Eukaryotic DNA Methylation and the Pursuit of Safer Sex. Current Biology 20, R780–R785. 10.1016/j.cub.2010.07.007

